# Xist RNA binds select autosomal genes and depends on Repeat B to regulate their expression

**DOI:** 10.1101/2024.07.23.604772

**Authors:** Shengze Yao, Yesu Jeon, Barry Kesner, Jeannie T Lee

## Abstract

Xist, a pivotal player in X chromosome inactivation (XCI), has long been perceived as a cis-acting long noncoding RNA that binds exclusively to the inactive X chromosome (Xi). However, Xist’s ability to diffuse under select circumstances has also been documented, leading us to suspect that Xist RNA may have targets and functions beyond the Xi. Here, using female mouse embryonic stem cells (ES) and mouse embryonic fibroblasts (MEF) as models, we demonstrate that Xist RNA indeed can localize beyond the Xi. However, its binding is limited to ∼100 genes in cells undergoing XCI (ES cells) and in post-XCI cells (MEFs). The target genes are diverse in function but are unified by their active chromatin status. Xist binds discretely to promoters of target genes in neighborhoods relatively depleted for Polycomb marks, contrasting with the broad, Polycomb-enriched domains reported for human XIST RNA. We find that Xist binding is associated with down-modulation of autosomal gene expression. However, unlike on the Xi, Xist binding does not lead to full silencing and also does not spread beyond the target gene. Over-expressing Xist in transgenic ES cells similarly leads to autosomal gene suppression, while deleting Xist’s Repeat B motif reduces autosomal binding and perturbs autosomal down-regulation. Furthermore, treating female ES cells with the Xist inhibitor, X1, leads to loss of autosomal suppression. Altogether, our findings reveal that Xist targets ∼100 genes beyond the Xi, identify Repeat B as a crucial domain for its in-trans function in mice, and indicate that autosomal targeting can be disrupted by a small molecule inhibitor.

## INTRODUCTION

The unequal sex chromosome composition between male (XY) and female (XX) placental mammals necessitates dosage compensation of the X chromosome ^1,2^. X chromosome inactivation (XCI) specifically evolved to ensure equal X-linked gene dosage between the sexes. During early embryogenesis, one of the two X chromosomes in female cells is randomly silenced, leading to a mosaic of cells in which X-linked genes can be expressed from either the maternal or paternal X chromosome ^3–5^. The long non-coding RNA Xist is an essential regulator of the XCI process ^6,7^. At the onset of XCI, Xist is expressed exclusively from the future inactive X chromosome (Xi) and then selectively spreads *in cis* along the chromosome to initiate formation of heterochromatin via association with chromatin-modifying complexes and alterations in 3D chromosome structure (reviewed in ^2,8–12^). From early RNA fluorescence in situ hybridization (FISH) experiments, it was shown at a cytological level that Xist binds only to the X-chromosome which transcribes the RNA ^6,13^. In *Xist* transgenesis studies, the RNA is also observed to localize exclusively in cis to the transgene, even on autosomes ^14–20^. In more recent years, technical advances in epigenomic mapping of RNA confirmed the early cytological data and provided a map of Xist binding sites in cis at kilobase resolution ^21,22^. Furthermore, genetic analysis of the X-inactivation center revealed a Xi-specific nucleation site for the initial binding of Xist that then enabled the RNA to spread exclusively *in cis* ^23^. Altogether, these studies solidified the view that Xist RNA is a cis-acting RNA.

However, studies have long documented the potential for Xist to spread beyond the X-chromosome under various non-physiological conditions. When Xist is overexpressed, the RNA can diffuse to bind neighboring chromosomes ^15,23,24^. Furthermore, when Xist’s nucleation site is mutated, Xist will diffuse and bind other chromosomes in trans ^23^. A recent study has also pointed to broad XIST binding patterns outside of the Xi in human naïve stem cells and suggested autosomal targets ^25^. These findings prompt interesting questions about the binding dynamics and properties of Xist on autosomes, its potential to spread locally in autosomal domains, and its impact on gene regulation and cellular physiology. To investigate, here we examine the capacity for Xist to spread beyond traditional boundaries. We use mouse embryonic stem cells (mESC) and mouse embryonic fibroblasts (MEF) in order to capture both the establishment and maintenance phases of XCI. Intriguingly, we identify about 100 binding sites on autosomes in cells undergoing XCI as well as post-XCI cells. We demonstrate discrete binding sites, rather than broad binding domains. Transcriptomic analysis reveals a selective downregulation, but not silencing, of associated genes. We also probe a requirement for Xist’s Repeat B (RepB) motif and the ability to perturb autosomal effects by treating cells with a small molecule inhibitor of Xist RNA.

## RESULTS

### Epigenomic mapping of Xist binding reveals autosomal targets in mouse cells

To explore the possibility of Xist extending its binding to autosomal sites beyond the X chromosome, we conducted CHART (Capture Hybridization of Associated RNA Targets) ^21, 26^ in female ES cells. To capture Xist RNA, we used oligoprobes antisense to Xist and then pulled down interacting chromatin for deep sequencing of the associated DNA. To control for any direct hybridization to DNA that could occur independently of Xist RNA, we conducted CHART in parallel using “sense” probes that should not hybridize to Xist. This helps us distinguish signals genuinely due to Xist RNA from those that could arise from non-specific probe binding to DNA. We note that some of the sense probes could hybridize to Tsix RNA, but Tsix expression is normally down-regulated by day 4 of ES differentiation ^27^. We also performed CHART using male ES cells, which do not upregulate Xist expression, to define the background level of binding. Notably, male ES cells express Tsix RNA at day 0 ^27^ and therefore also assist in excluding non-Xist RNA binding during analysis. Together the male and sense controls allowed us to account for non-specific interactions and background noise. In addition to these controls, we performed 2 biological replicates and analyzed only overlapping signals between the 2 replicates to ensure the reliability and reproducibility of our results. The Pearson and Spearman correlation analysis showed good reproducibility between replicates (Figure S1A). We also normalized to input DNA to account for differences in chromatin preparation and sequencing depth.

Xist RNA is transcriptionally upregulated during female ES cell differentiation when XCI is induced. To profile the dynamics during cell differentiation and XCI, we examined wild-type (WT) female ES cells in the undifferentiated state (day 0) and at three differentiated stages (day 4, 7, and 14). Antisense oligonucleotides targeting Xist efficiently pulled down Xist RNA and associated chromatin targets, in excess of the “background” observed with the sense probe, and male control (Figure S1B). Principal component analysis (PCA) of CHART-seq data from day 4 and Xist coverage on X-linked genes revealed that Xist CHART signals in female cells were distinct from those of sense probe and male controls (Figure S1C). This PCA separation indicates a clear difference in Xist binding in females as compared to controls. As expected, the Xist CHART signal in WT female ES cells displayed strong binding at the Xist locus and genes subject to XCI, such as *Cdkl5* and *Mecp2*, while showing significantly weaker binding at the escapee gene, *Kdm6a*. In contrast, CHART experiments using sense probes and male ES cells, which lack Xist expression, showed minimal binding to these genes (Figure S2A-S2D, S3). Because Xist spreads *in cis* along the Xi to cover much of the chromosome, we first examined total Xist coverage (normalized to input), rather than calling peaks ^28^. Day 0 ES cells showed no enrichment for Xist binding, consistent with Xist being in the uninduced state (Figure 1A). Upon cell differentiation, enriched binding of Xist was consistently observed across all time points (Figure 1A). Greatest enrichment on the X chromosome was observed on day 7 (Figure 1B), in agreement with de novo XCI when Xist is needed at highest concentration ^21,29^.

**Figure 1.**
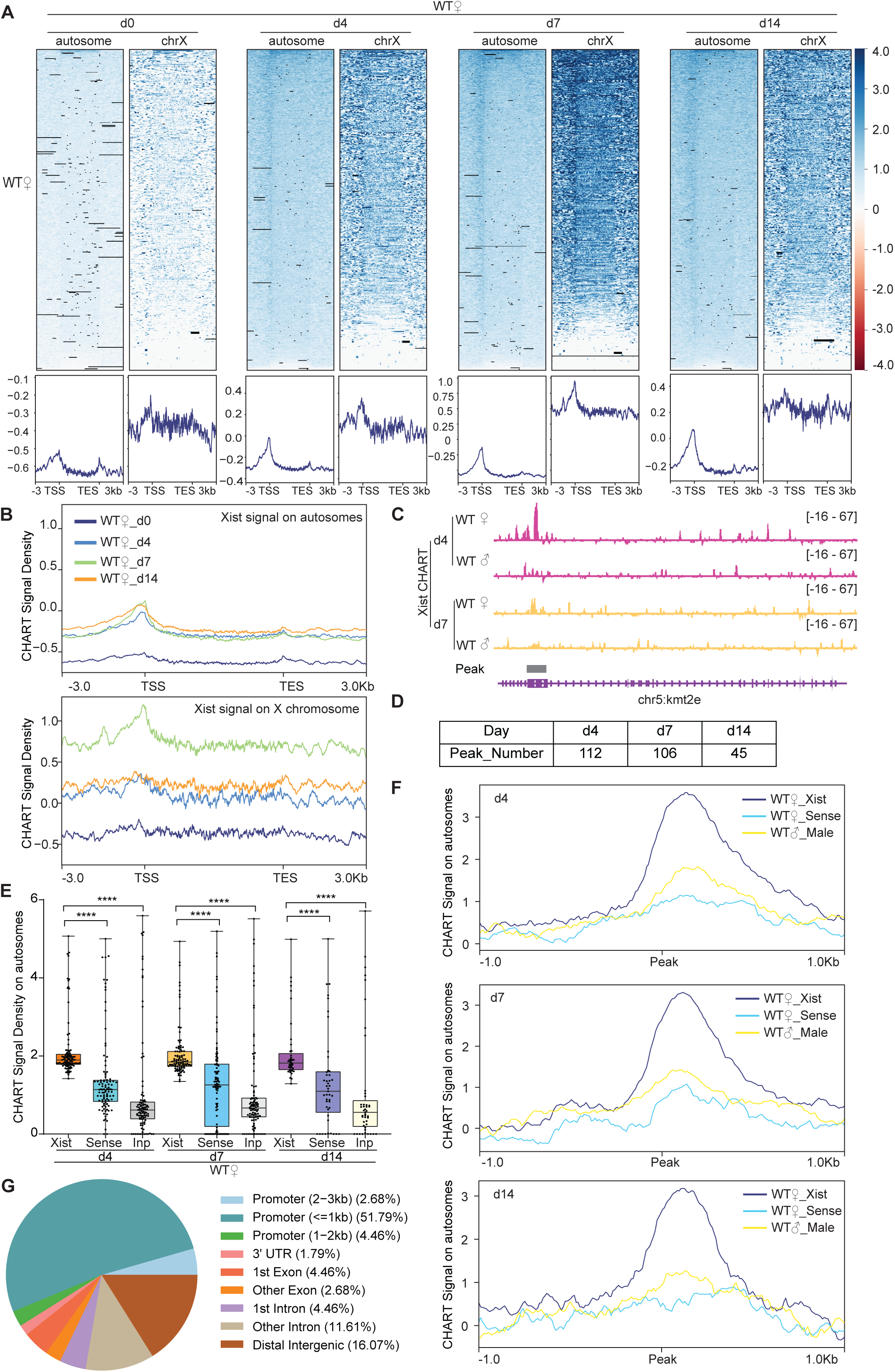
CHART-Seq reveals ∼100 discrete binding sites in autosomal genes. **A.** Top: Heatmaps of Xist coverage on autosomes and chromosome X genes in WT female ES cells at day 0, day 4, day 7, and day 14. Bottom: Average profiles shown as metagene maps of the data in the top panel. TSS, transcription start site. TES, transcription end site. **B.** Average profile of Xist coverage on autosomes and X chromosome genes in WT female ES cells at day 0, day 4, day 7, and day 14. **C.** Representative site-specific binding of Xist on autosome locus (*Kmt2e*) in WT female ES cells at day 4 and day 7, WT male ES cells are used as control. **D.** Number of Xist peaks determined by MACS2 peak calling on autosomes in WT female ES cells at day 4, day 7, and day 14. **E.** Xist CHART signal coverage on Xist-autosomal peak region (top 100 peaks) in WT female ES cells at day 4, day 7, and day 14. Sense and input are used as control. P-values are determined using the Wilcoxon rank sum test. **F.** Average profile of CHART (subtracted input) signal on Xist-autosomal peak region (top 100 peaks) in WT female and male ES cells at day 4, day 7, and day 14. Sense and WT male ES are used as control. **G.** Feature annotation of Xist binding loci on autosomes by ChIPseeker in WT female ES cells at day 4.

Intriguingly, on day 7, only ∼10% of the reads mapped to the X chromosome (Figure S1B), prompting the question of whether Xist might have non-X-linked targets as well. Indeed, although the X chromosome was most enriched for Xist, we noticed that autosomes also showed highly reproducible peaks, beginning at day 4 and increasing across the differentiation time points (Figure 1A). In contrast to broad binding pattern covering the entire X chromosome (Figure S2E), Xist binding patterns on autosomes trended towards sharp peaks (Figure 1C and S2F). To determine if the peaks were statistically significant, we called Xist peaks using MACS2 on the Xist CHART data. We restricted the analysis to autosomal reads to increase the sensitivity of the analysis. We performed peak-calling separately for each CHART biological replicate. Irreproducible Discovery Rate (IDR) analysis indicated a strong correlation between the two replicates (Figure S4A). We used *bedtools intersect* to identify significant peaks that overlapped between the two replicates and only used overlapping peaks in subsequent analyses. Overall, Xist coverages in significant peaks of female cells were substantially greater than in the negative controls (Figure 1E-1F).

Intriguingly, we found hundreds of significant peaks across autosomes between days 4-14 of differentiation (Figure 1D, S4B, Tables S1-S6). Notably, Xist binding sites on autosomes were disproportionately located in promoter regions (Figure 1A and 1G), hinting at a potential role for Xist in autosomal gene regulation. We conclude that, in addition to the Xi of differentiating ES cells, mouse Xist RNA selectively binds ∼100 autosomal targets, preferentially at promoter regions. This promoter-dominant profile contrasts with XIST patterns identified in human cells, which tend to demonstrate broad regions of coverage over genes ^25^. Although the Gene Ontology (GO) analysis of these Xist autosomal target genes did not yield statistically significant enrichment for specific biological processes or pathways, a closer examination reveals that these genes have diverse functions. They are involved in processes such as cancer development (*Bcl7b*), POLII transcription regulation (*Med16*), amino acid transport (*Slc36a4*), RNA binding (*Rbm14* and *Stau2*), among other processes. Together with previously published work in human cells ^25^, our findings suggest that Xist may have a broad regulatory impact on a variety of autosomal genes, extending its influence beyond X-chromosome inactivation.

### Deleting RepB causes a loss of binding to autosomal target genes

Prior work showed that Xist’s Repeat B (RepB) element is essential for Polycomb recruitment ^30–33^. RepB is also required for proper spreading and localization of Xist RNA to the Xi: Without RepB, the Xist RNA cloud was observed to adopt a dispersed appearance consistent with diffusion of the RNA away from the Xi ^32^. Given the partial loss of attachment to the Xi, here we asked if the Xist diffusion could lead to enhanced binding to autosomal targets. We conducted CHART experiments in RepB deletion (ΔRepB) female ES cells at various differentiation stages and quantitated the degree of X-linked versus autosomal binding (Figure 2). Although Xist retained partial binding to the X-chromosome, there was a significant decrease in localization along X-linked genes along all differentiation days, especially at day 7 (Figure 2A), consistent with RepB being essential for attachment of Xist to the Xi ^32^. Intriguingly, however, the loss of Xi binding was not accompanied by any significant increase in Xist coverage at the same autosomal targets, at any differentiation day (Figure 2A). In fact, there appeared to be decreased binding to the ∼100 autosomal genes (Figures 2E-2F, S4C-S4D). However, there was not a change up- or down-wards in the number of autosomal targets (Figures 2B-2C, S4B, S4E, Tables S1-S6). A similar number of Xist peaks across autosomes in ΔRepB cells was observed and the autosomal targets remained similar. Moreover, in the absence of RepB, Xist retained a preference to bind promoter regions (Figure 2D). Thus, RepB is required not only for Xist to localize to the X-chromosome but also for its localization to the ∼100 autosomal genes.

**Figure 2.**
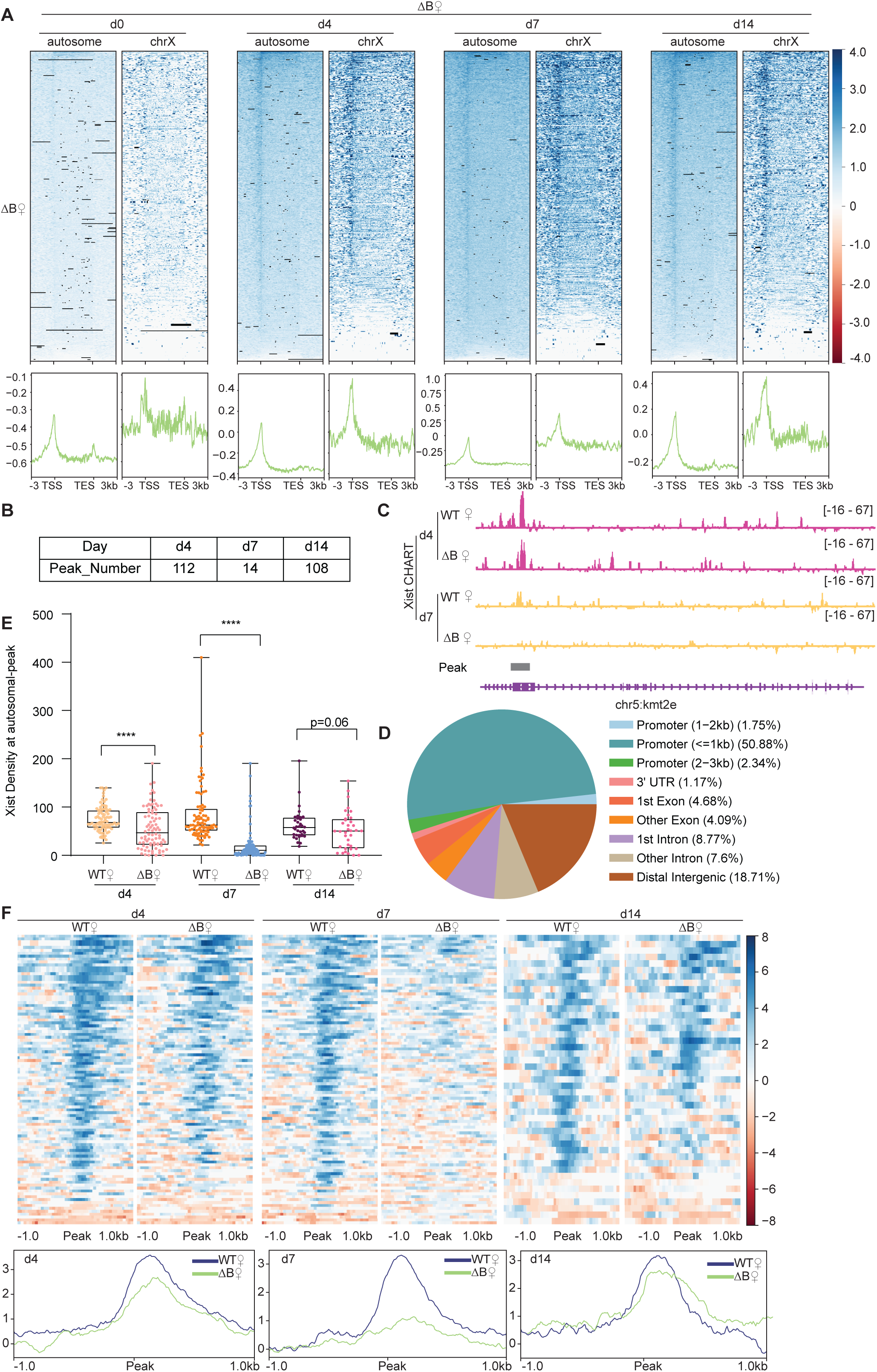
Xist’s Repeat B is required for proper binding of Xist to autosomal genes. **A.** Top: Heatmaps of Xist coverage on autosomes and chromosome X genes in WT female ES cells at day 0, day 4, day 7, and day 14. Bottom: Average profiles shown as metagene maps of the data in the top panel. **B.** Number of Xist peaks determined by MACS2 peak calling on autosomes in ΔRepB female ES cells at day 4, day 7, and day 14. **C.** Representative site-specific binding of Xist on autosome locus (*Kmt2e*) in WT and ΔRepB female ES cells at day 4 and day 7. **D.** Feature annotation of Xist binding loci on autosomes by ChIPseeker in ΔRepB female ES cells at day 4. **E.** Xist CHART signal coverage on Xist-autosomal peak region in WT and ΔRepB female ES cells at day 4, day 7, and day 14. P-values are determined using the Wilcoxon rank sum test. **F.** Top: Heatmaps of Xist coverage on Xist-autosomal peak region (top 100 peaks) in WT and ΔRepB female ES cells at day4, day 7, and day 14. Bottom: Average profiles shown as metagene maps of the data in the top panel.

### Xist RNA down-regulates autosomal genes in a RepB-dependent manner

The promoter-specific binding pattern of Xist on autosomal targets contrasts sharply with the broad binding pattern of Xist on the Xi (Figure 1A) and with broad regions identified for human autosomal genes ^25^. Given the preference for autosomal promoters, we investigated whether Xist modulates expression of the associated autosomal genes. To address this question, we conducted transcriptome sequencing on WT female ES cells across day 0, 4, 7, and 14, and compared the profile to those of ΔRepB female ES cells. The RNA-seq data showed that Xist expression increased during the establishment phase of XCI in differentiating ES cell, reaching its peak at day 7 (Figure S5A), consistent with super-resolution data indicating that Xist is present at ∼300 copies/cell during de novo XCI relative to the ∼100 copies/cell during the maintenance phase ^29^. Xist expression followed a similar dynamic in ΔRepB female cells (Fig. S5A). However, deletion of Xist’s RepB resulted in increased X-linked genes expression in differentiating female ES cells (Figures 3D and S5B-5C), consistent with a previous report ^32^. To analyze the characteristics of genes bound by Xist on autosomes, we categorized all refSeq genes based on their expression levels at each differentiation day. The genes were divided into 5 quintiles (Q1 silent, Q2 low, Q3 moderate, Q4 high, Q5 highest). Consistent with Xist’s behavior on the X-chromosome ^21,22^. Xist also favored binding to actively expressed genes on autosomes (Figure 3A). To examine gene expression levels as a function of distance from the Xist binding site, we divided neighboring genes into 10- to 100-kb bins and observed that genes at <10 kb showed highest Xist levels (Figure 3B, Figure S6A-S6B). These findings support Xist’s favoring actively expressed genes. We then examined Polycomb marks in relation to the autosomal Xist domains. In agreement with the Xist’s predilection for active genes, we observed lower coverage of Polycomb marks associated with Xist RNA^12,17,30–34^, H3K27me3 and H2AK119ub (associated with PRC2 and PRC1 respectively), as shown by lower coverages within the 10-kb versus 50-kb neighborhoods (Figure 3C, Figure S6C). These findings also contrasted with observations in human cells, where PRC2/H3K27me3-enriched regions were favored ^25^. Our findings indicate that WT Xist transcripts favor binding to a select set of actively transcribed autosomal genes and these genes are not marked by Polycomb in mouse cells.

**Figure 3.**
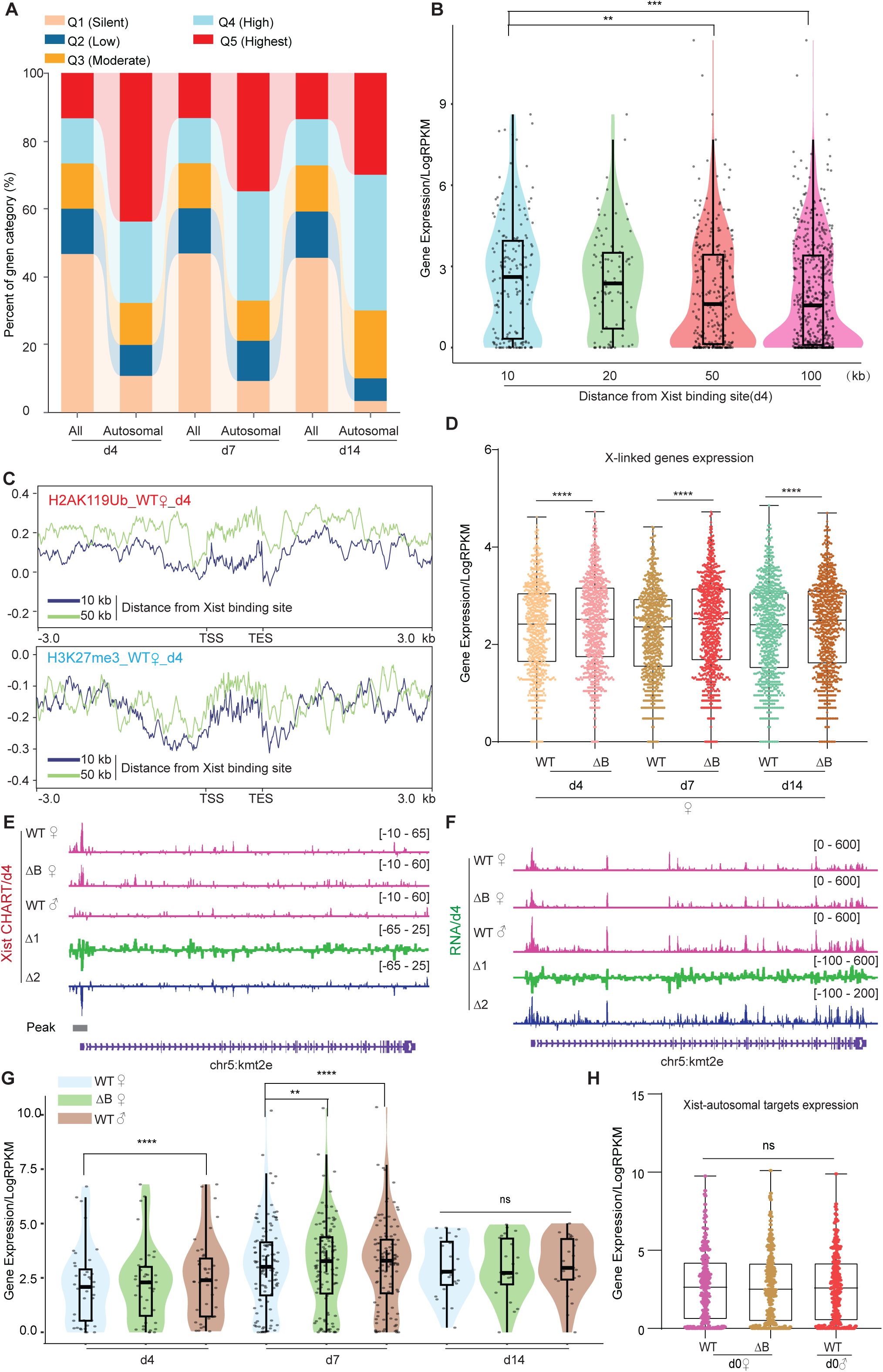
Autosomal target genes are actively transcribed. **A.** Xist preferentially targets actively expressed genes. The proportion of overall genes (label as “All”) and Xist targets on autosomes expression levels (label as “Autosomal”) in different expression categories. Genes are classified into distinct categories (Q1-Q5) based on RPKM values, representing various levels of expression. Category Q1 includes non-expressed genes (RPKM=0), while categories Q2, Q3, Q4, and Q5 represent 25%, 50%, 75%, and 100% expression, respectively. **B.** Genes bound by Xist exhibit elevated expression levels compared to surrounding regions. The analysis of gene expression within the 10, 20, 50, and 100-kilobase binding regions of Xist is conducted in WT female ES cells at day 4. P-values are determined using the Wilcoxon rank sum test. **C.** The H2AK119ub and H3K27me3 (Chip-Seq) average profile of genes within the 10 and 50-kilobase binding regions of Xist in WT female cells at day 4. **D.** Evaluation of gene expression levels for XCI genes in WT and ΔRepB female ES cells at day4, day7, and day 14. P-values are determined using the Wilcoxon rank sum test. **E-F.** Representative CHART-seq (F) and RNA-Seq (G) patterns of an autosomal gene bound by Xist *(Kmt2e*) at day 4. Change in coverage (Δ1 and Δ2) is shown below (Δ1 for ΔRepB♀ -WT♀, and Δ2 for WT♂ - WT♀). **G.** Assessing gene expression levels of Xist targets on autosomes (10 kb within the peak region) in WT, ΔRepB female ES cells, and male ES cells at different time points. P-values are determined using the Wilcoxon rank sum test. **H.** Gene expression levels for Xist targets on autosomes (identified in day4 and day7) in undifferentiated WT, ΔRepB, female and male ES cells (day 0) show no obvious changes. 2 biological replicates were used. P-values are determined using the Wilcoxon rank sum test.

Given the association with active autosomal genes, we next asked whether Xist’s binding modulates their expression. We performed RNA-seq analysis in two biological replicates in the ES differentiation time series. First, we profiled gene expression in female ES cells relative to male ES cells, reasoning that the lack of Xist RNA in male ES cells might lead to a difference in female versus male expression of autosomal target genes. Indeed, among the ∼100 target genes, there was significantly greater expression in male cells (Figure 3E-3G), suggesting that Xist might also negatively regulate the select genes on autosomes. To test this idea, we took advantage of the ΔRepB mutation, as RepB is required for proper silencing of Xi genes ^32^. Transcriptomic analysis demonstrated that perturbing RepB resulted in loss of repression of autosomal targets. A detailed examination focusing on the top 100 peaks of Xist-bound genes on autosomes unveiled a down-modulation especially evident on days 4 and 7 of ES cell differentiation (Figure 3G and S7A-S7D). The Xist-bound autosomal genes on day 14 of ES cell differentiation like *Ces1l* or non-targets of Xist on autosomes such as *Kmt2c* did not demonstrate significant changes in gene expression (Figure S7E and S7F). We note, however, that unlike on the Xi, autosomal binding of Xist did not lead to full silencing of the target gene. Rather, the effect was a partial repression. In ΔRepB cells, reduced autosomal binding resulted in a blunting of this repression (Figure 3F-3G and S7A-S7D). On the other hand, at day 0 when Xist had yet to be upregulated, ΔRepB female and male ES cells showed no significant expression changes at Xist-autosomal targets (Figure 3H). To confirm that the observed changes in autosomal genes are due to Xist binding, we performed a similar analysis on 100 randomly selected autosomal genes that did not have Xist binding (“non-targets”). The non-targets did not show preference for specific autosomal gene region (Figure S8A-C) and showed no significant difference in Polycom enrichment (Figure S8D) or gene expression (Figure S8E). These results demonstrate that autosomal binding of Xist RNA directly leads to a suppression of gene expression in a RepB-dependent manner.

### Effects of Xist overexpression and X1 drug treatment on autosomal genes

To seek further evidence for autosomal effects of Xist binding, we conducted two orthogonal lines of experimentation. First, we asked the reciprocal question and determined whether ectopically overexpressing Xist leads to increased autosomal repression. We examined two previously published transgenic *Xist* cell lines, Tg_1 and Tg_2 ^35^, to test whether — *<u>in the hands of other investigators</u>* — cell lines expressing Xist also demonstrated autosomal targeting (Figure 4A). These two transgenic (Tg) females cell lines respond to doxycycline to induce Xist expression on autosomes (specifically chromosome 12, with two different transgene insertion sites). Transcriptomic analysis of these two Tg mouse ES cells upon neuronal differentiation showed a significant suppression of X-linked genes (e.g., *Med14*) in comparison to doxycycline-treated wildtype cells (Figure 4B and S9A). Autosomal Xist targets, as exemplified by *Bcl7b* and *Rbm14*, were also significantly suppressed beyond what was observed in WT ES cells (Figure 4C and S9B). In contrast, non-targets of Xist such as *Stau1* did not demonstrate significant changes in gene expression (Figure 4E and 4F). Looking across all autosomal target genes, we observed a significant decrease in mean expression in the Xist overexpressing cell lines (Figure 4D), further implicating Xist binding in direct regulation of autosomal gene targets. The fact that the autosomal changes were also observed in datasets generated by other investigators strengthen our conclusions.

**Figure 4.**
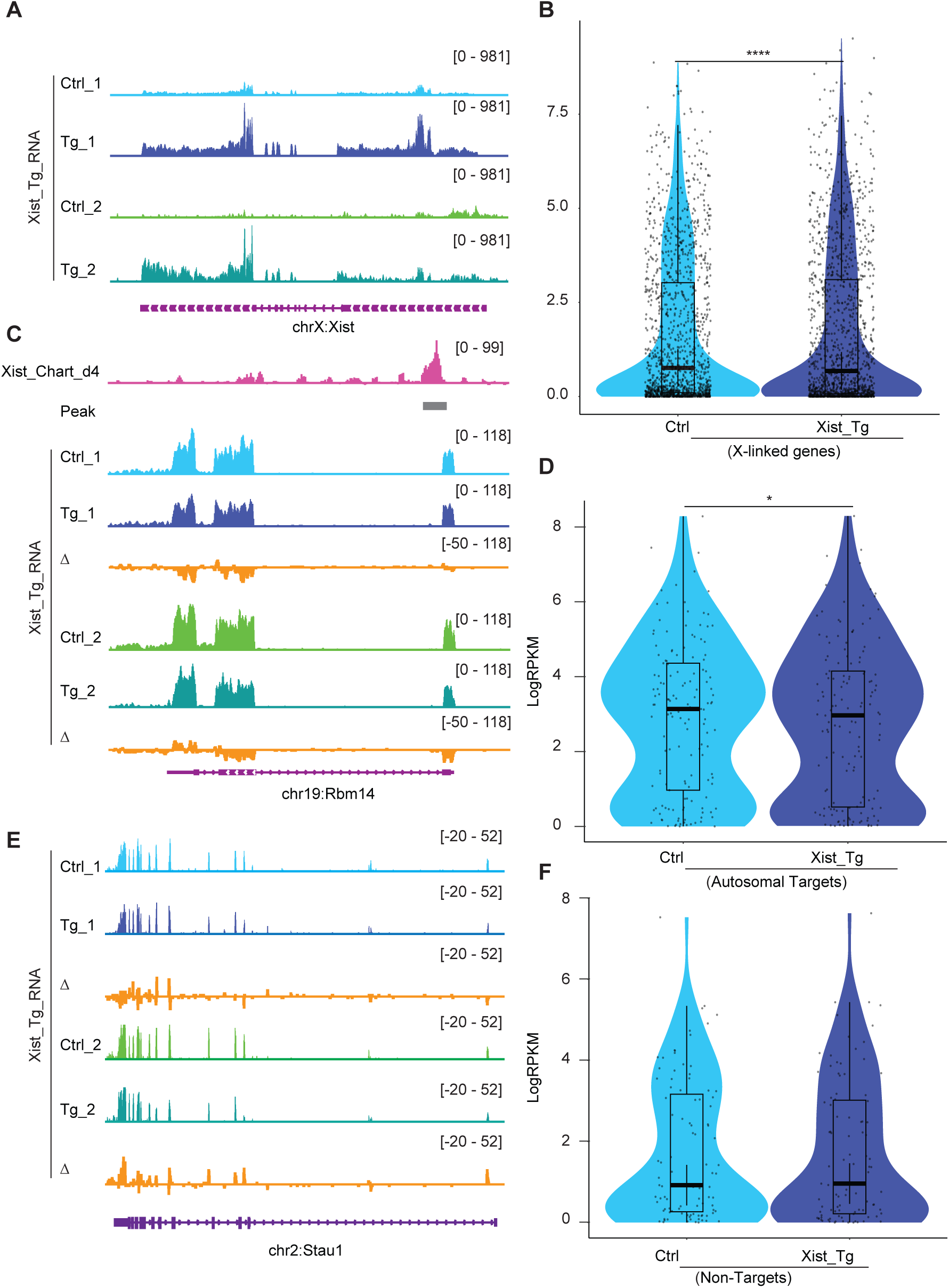
Xist binding suppresses, but does not silence, autosomal gene expression. **A.** RNA-seq track shows the Xist expression level in ectopic doxycycline responsive Xist overexpressed ES cells (129/Sv-Cast/Ei) of neuronal differentiation (Tg-1 and Tg-2 have different insertion sites), the control (Ctrl) was doxycycline-treated wildtype cell. **B.** Analysis of gene expression levels for XCI genes in Xist Tg ES cells (129/Sv-Cast/Ei) at of neuronal differentiation. P-values are determined using the Wilcoxon rank sum test. **C.** RNA-seq track illustrating expression alongside representative autosome genes with Xist binding (*Rbm14*) in Xist Tg ES cells (129/Sv-Cast/Ei) of neuronal differentiation. Change in coverage (Δ) is shown below (Tg - Ctrl). **D.** Analysis of gene expression levels for Xist autosomal targets in Xist Tg ES cells (129/Sv-Cast/Ei) of neuronal differentiation. P-values are determined using the Wilcoxon rank sum test. **E.** RNA-seq track illustrating expression level of autosome genes without Xist binding (*Stau1*) in Xist Tg ES cells (129/Sv-Cast/Ei) of neuronal differentiation. Change in coverage (Δ) is shown below (Tg - Ctrl). **F.** Analysis of gene expression levels for Xist non-targets on autosomes in Xist Tg ES cells (129/Sv-Cast/Ei) of neuronal differentiation. P-values are determined using the Wilcoxon rank sum test.

In the second line of investigation, we deployed X1, a small molecule inhibitor of Xist that was previously shown to block initiation of XCI in female ES cells through Xist’s Repeat A (RepA) motif ^36^. We reasoned that if the mechanism of autosomal gene suppression were similar to XCI, treating cells with X1 could also blunt autosomal gene suppression. Indeed, just as X-linked genes such as *Mecp2* failed to undergo silencing after 5 days of differentiation, transcriptomic analysis showed that autosomal gene targets also failed to be suppressed in the presence of X1 (Figure 5A-C). On average, there was a significant upregulation of autosomal targets on day 5 of differentiation and X1 treatment (Figure 5E). Again, non-targets of Xist such as *Cbx2* did not demonstrate significant changes in gene expression (Figure 5D and 5F). Analysis of ChIP-seq data for H3K27me3 indicated a reduction of the PRC2 mark within the Xist target gene and flanking regions (Figure 5G and 5H), consistent with X1 blocking RepA’s recruitment of PRC2 ^36^. In contrast, Xist non-target genes show no significant changes in PRC2 signal (Figure 5I). These additional perturbation studies reinforce the notion that autosomal binding of Xist RNA directly suppresses autosomal targets. Thus, it is possible to disrupt autosomal Xist binding by administering a small molecule inhibitor of Xist.

**Figure 5.**
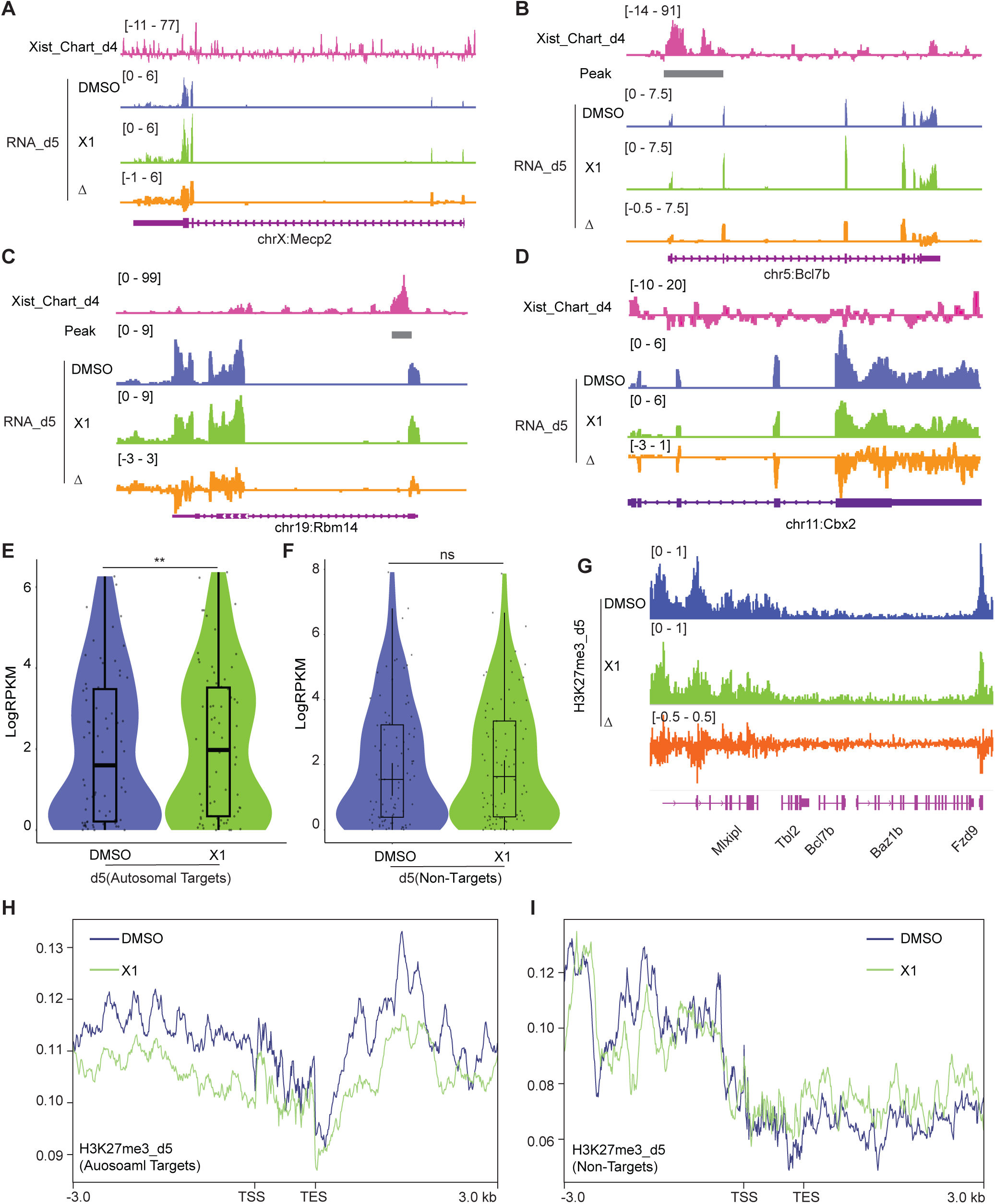
Treating cells with the X1 inhibitor of Xist RNA perturbs autosomal target genes. **A-D.** RNA-seq track depicting *Mecp2* (A) (an X-linked gene), *Bcl7b* (B) and *Rbm14* (C) (Xist-regulated autosome target), and *Cbx2* (D) (autosomal gene without Xist binding) expression in DMSO (control) and Xist inhibitor (X1) treated differentiated WT female ES cells at day 5 of differentiation. Change in coverage (Δ) is shown below (X1 - DMSO). The Xist peak on autosomes is from CHART at day 4. **E.** Evaluation of gene expression levels for Xist autosomal targets in DMSO (control) and X1 treated WT female ES cells at day 5 of differentiation. The Xist peak on autosomes is from CHART at day 4. P-values are determined using the Wilcoxon rank sum test. **F.** Evaluation of gene expression levels for Xist non-targets on autosomes in in DMSO (control) and X1 treated WT female ES cells at day 5 of differentiation. P-values are determined using the Wilcoxon rank sum test. **G.** ChIP-seq track for H3K27me3 on *Bcl7b* (an Xist autosomal target, within a 160 kb window) in DMSO (control) and X1 treated WT female ES cells at day 5 of differentiation. Change in coverage (Δ) is shown below (X1 - DMSO). **H-I.** Profile plot displaying H3K27me3 levels for Xist-autosomal targets flanking genes (H) (within a 100 kb window) or Xist non-targets (I) in DMSO (control) and X1 treated WT female ES cells at day 5 of differentiation. The Xist autosomal target is from CHART at day 4.

### Xist RNA also binds to autosomal genes in post-XCI cells

Lastly, because the establishment and maintenance phases of XCI show differential sensitivity to loss of Xist ^37–40^, we asked if autosomal repression exhibits a similar sensitivity. To address this, we turned to mouse embryonic fibroblasts (MEF), where XCI is well established. We first conducted CHART assays to determine whether Xist binds to the Xi and autosomes in a similar way in MEFs (Figure S10A-10B). Because the autosomal signals displayed peak-like characteristics, we used MACS software to call significant peaks on autosomes to determine if there was an increase in promoter-specific binding of the autosomal genes. We observed that, while Xist continued to favor targeting promoter regions in both WT and ΔRepB MEFs (Figure 6A-6C), Xist peaks did not increase in size in ΔRepB MEFs; rather, the peaks decreased in size (Figure 6A-6C) — a result similar to that in ΔRepB ES cells (Figure 2). Furthermore, there was a considerable decrease in the number of significant peaks in ΔRepB MEFs when compared to WT MEFs (Figure 6A). In WT MEFs, Xist clearly favored Xi binding, but Xist signals could also be visualized on autosomes (Table S7-S10, Figure 6E), in agreement with the results in ES cells (Figure 1). In ΔRepB MEFs, there was a partial loss of Xi localization, aligning with previous findings ^32^ (Figure 6D-6F). Consistent with these findings, RNA-seq analysis of ΔRepB MEFs did not reveal any significant change in expression of Xi genes (e.g., *Mecp2*) and autosomal target genes (e.g., *Rbm14*) (Figure 7A-7D).

**Figure 6.**
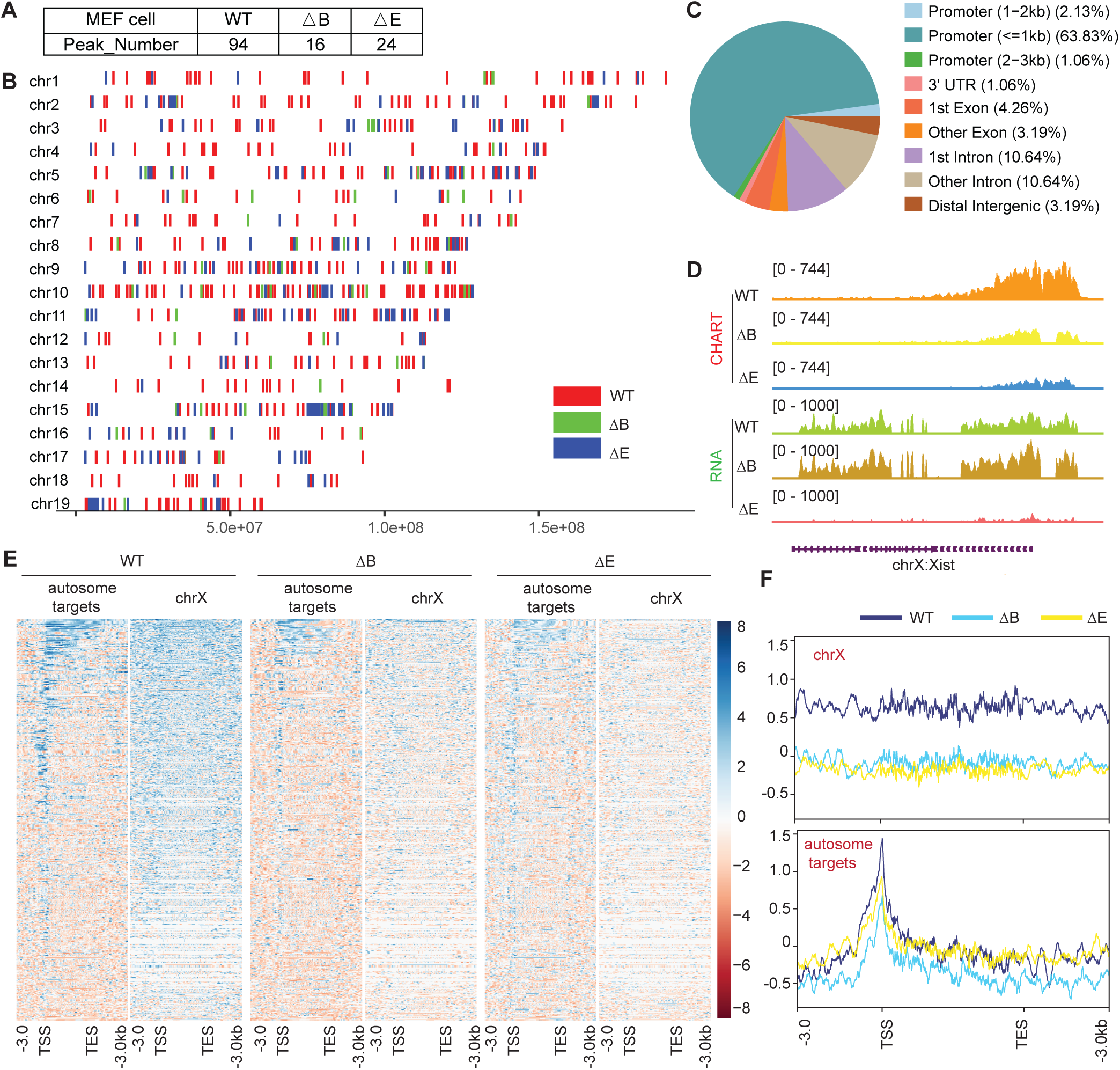
Xist RNA also binds autosomal genes in post-XCI MEF cells. **A.** Xist peak number (MACS2 peak calling) on autosomes in WT, ΔRepB, and ΔRepE female MEF cells. **B.** Xist peak patterns (MACS2 peak calling) on autosomes in WT, ΔRepB, and ΔRepE female MEF cells. **C.** Feature annotation of Xist binding loci on autosomes in WT female ES cells. **D.** Representative CHART-seq and RNA-seq track patterns of Xist in WT, ΔRepB, and ΔRepE female MEF cells. **E-F.** Heatmaps (E) and average profiles (F) depicting Xist coverage on autosome targets and chromosome X in WT, ΔRepB, and ΔRepE female MEF cells.

**Figure 7.**
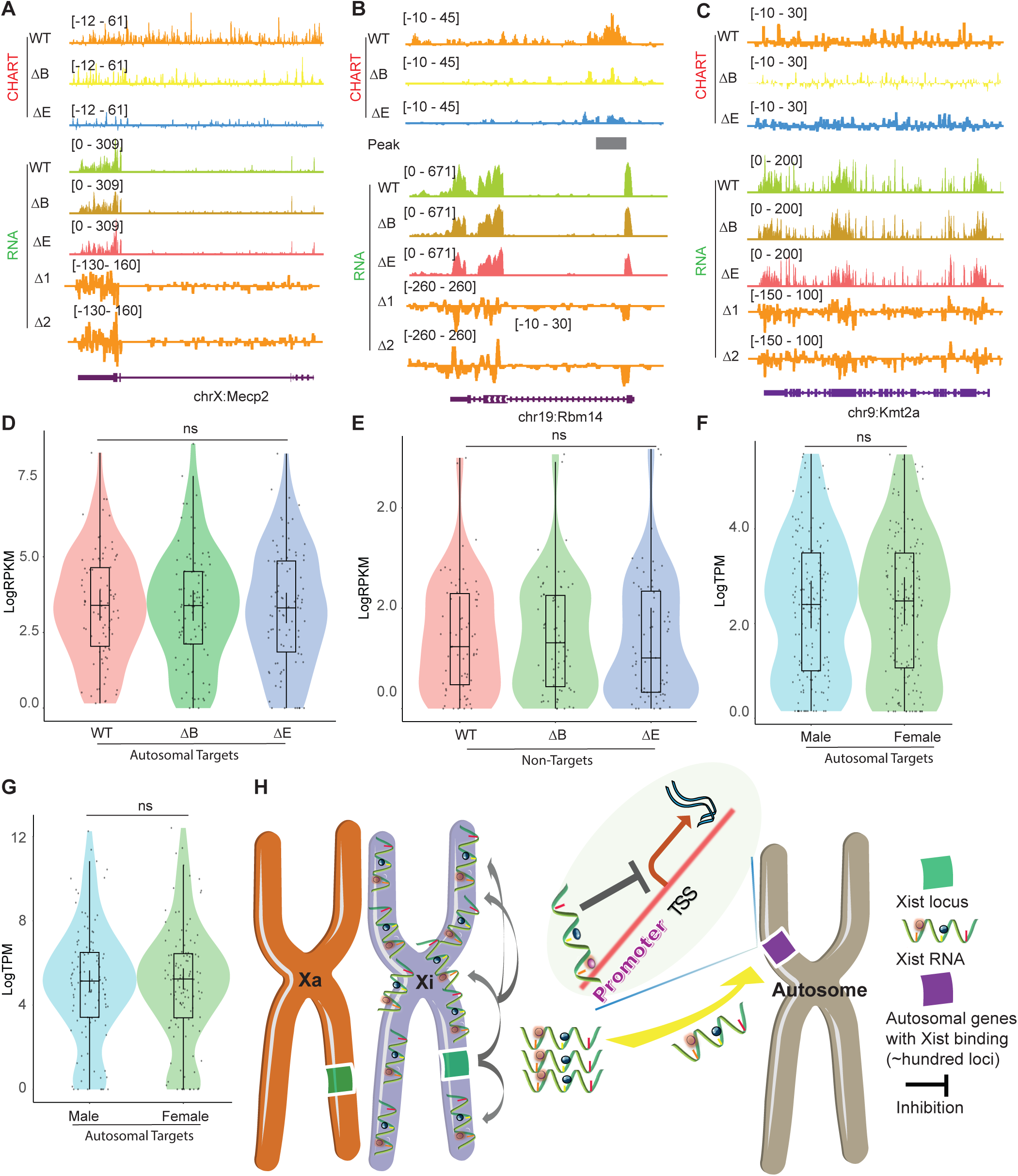
Xist autosomal binding does not alter gene expression in MEF Cells. **A-C.** Representative CHART-seq and RNA-seq track patterns of Xist autosomal binding genes such as *Mecp2* (A) and *Rbm14* (B), and autosome genes without Xist binding such as *kmt2a* (C) in WT, ΔRepB, and ΔRepE female MEF cells. Change in coverage (Δ1 and Δ2) is shown below (Δ1 for ΔRepB -WT, and Δ2 for ΔRepE -WT♀). **D-E.** Gene expression levels for Xist targets (D) or non-targets (E) on autosomes in WT, ΔRepB, and ΔRepE female MEF cells show no obvious changes. P-values are determined using the Wilcoxon rank sum test. **F-G.** Gene expression levels for Xist targets on autosomes in male and female MEF cells show no obvious changes. P-values are determined using the Wilcoxon rank sum test. **H.** Schematic of the Xist autosome binding pattern influences the gene expression. During the differentiation process of female ES cells, the Xist is specifically expressed from the Xist loci of one X chromosome (Xi). It then binds to the Xist loci of the Xi *in cis*, spreading across this chromosome and effectively silencing the expression of most of its genes. Additionally, some Xist complex will also bind to hundreds of loci on autosomes, thereby inhibiting the expression levels of target genes located there.

We then asked if the retention of some Xist binding on the Xi could explain the lack of transcriptomic difference in ΔRepB MEFs. To investigate, we utilized a deletion of Xist’s Repeat E (ΔRepE), which was previously demonstrated to severely abrogate localization of Xist to the Xi ^41,42^. We reasoned that the severe loss of Xist binding might unmask a transcriptomic difference. As expected, we observed that Xist signals were somewhat more reduced on the Xi in ΔRepE MEFs compared to ΔRepB cells (Figure 6E-6F). Despite this reduction, peak coverages in autosomal target genes did not increase in ΔRepE MEFs (Figure 6E-6F). However, there was an overall decrease in the number of significant autosomal peaks in ΔRepE MEFs relative to WT cells (Figure 6A). Regardless, we observed no significant transcriptomic differences in ΔRepE MEFs relative to WT MEFs (Figure 7A-7E). Additionally, further examination of RNA sequencing data from male and female MEF cells in two published studies ^43,44^ corroborated that the expression levels of these autosomal Xist targets did not exhibit significant changes (Figure 7F and 7G). Altogether, the analysis in MEFs demonstrates that Xist continues to bind autosomal genes in post-XCI somatic cells. However, autosomal binding of Xist in post-XCI cells does not overtly impact expression of the associated autosomal genes. Nonetheless, we cannot exclude more subtle changes that do not meet the significance cut-off.

## DISCUSSION

Together with a backdrop of studies on ability of Xist RNA to diffuse and bind chromatin in trans ^15,23–25^, the results of our current work challenges the conventional narrative that Xist operates exclusively *in cis* on the Xi. We find that autosomal Xist targets are not numerous, possibly limited to only ∼100 in both pluripotent stem cells and in somatic cells. On autosomes, Xist does not spread and instead covers only a narrow region corresponding to single genes. The genes tend to be active genes, with Xist specifically targeting the promoters of those genes. Genetic analysis coupled to transcriptomic analysis showed that Xist down-regulates the target autosomal genes without silencing them. This effect leads to clear sex difference — where female cells express the ∼100 or so autosomal genes at a lower level than male cells in the mouse ES cell differentiation (Figure 7H). Thus, our findings redefine the scope of Xist’s functional repertoire and provide insights into the broader landscape of epigenetic regulation during cellular differentiation. Xist RNA therefore plays a more complex role than previously envisaged, with several implications and caveats.

First, the observed consistent binding of Xist to both the X chromosome and autosomes in female ES cells prompted further exploration of the intricate dynamics of Xist during cellular differentiation. Our categorization of genes based on their expression levels revealed a compelling correlation between Xist binding on autosomes and the expression levels of associated genes. Notably, Xist exhibited a preference for binding to active gene regions, as evidenced by the significantly higher expression of genes within the 10-kb range of Xist binding regions. The parallel upregulation of X-linked and autosomal genes in ΔRepB and male ES cells suggests a regulatory role of Xist in gene expression beyond its canonical function on the X chromosome. Therefore, the precise temporal modulation of Xist binding merits further investigation to elucidate its regulatory significance and during distinct stages of cell differentiation or development, particularly in understanding its autosomal targets which could have implications for gene regulation and cellular function.

Second, a recent study using an alternative Xist pulldown method, RAP-seq, also revealed the capability of Xist to bind to autosomes in naive human pluripotent stem cells (naive hPSCs) and mediate gene regulation ^25^. While we found that, on autosomes, Xist does not spread and instead covers only a narrow region corresponding to single genes, the previous study demonstrated broader regions of binding ^25^. Furthermore, our CHART experiments demonstrate Xist binding to autosomes in post-XCI cells (e.g., MEF), whereas the previous study found no significant binding beyond the stem cell stage ^25^. Our analysis also indicates that Xist’s autosomal targets tend to be Polycomb-depleted relative to neighboring genes, whereas the prior human study observed a positive correlation between XIST RNA and PRC2 marks. These discrepancies may arise from differences in the Xist pulldown methods (CHART versus RAP) employed by the two studies or from inherent differences between mouse and human systems.

Third, we considered the possibility that the binding of Xist to autosomes could merely be a consequence of Xist diffusion following saturation of binding sites on the Xi, rather than any programmed event during development. We are inclined to reject this notion and propose that Xist binding to autosomes is specifically programmed. Upon deleting RepB, the binding of Xist to the X chromosome weakens, but concomitantly, its binding to autosomal targets also diminishes (Figure 2). This suggests that Xist binding to autosomes is contingent upon Repeat B and is deliberate, rather than due to random Xist diffusion alone (in which case we would have expected increased autosomal binding). Additionally, following treatment with the Xist inhibitor X1, an increase in the expression of autosomal targets is observed (Figure 5), implying that the regulation of autosomes by Xist is not merely an overflow effect of Xist saturating the X chromosome.

In summary, our study advances understanding of Xist-mediated epigenetic regulation by highlighting unexplored interactions with autosomes. The identified correlations between Xist binding and gene expression involvement pose intriguing questions regarding the regulatory mechanisms governing these processes. These insights contribute to the evolving paradigm of Xist biology, underscoring the need for continued exploration of the complexity of epigenetic control mechanisms. Future investigations will delve deeper into the functional consequences of Xist binding on autosomes and explore the potential downstream effects on cellular differentiation, development, and diseases associated with its dysregulation, such as cancer, immunity, and neuron development ^25,45–49^. A comprehensive understanding of Xist’s influence beyond X chromosomes is crucial, particularly laying the groundwork for future exploration of Xist as a potential therapeutic target.

## METHODS

### Cell lines

The wild-type and Xist’s Repeat B or E deletion MEF cell lines (M. musculus/M. castaneus F1 hybrid) has been previously described and generated ^32^. Additionally, male Xist transgenic (Xist TG) MEF cells, previously denoted as “♂X+P” ^23^ were included in the study. MEFs were cultured in medium comprising DMEM, high glucose, GlutaMAX™ Supplement, pyruvate (Thermo Fisher Scientific), 10% FBS (Sigma), 25 mM HEPES pH 7.2-7.5 (Thermo Fisher Scientific), 1x MEM non-essential amino acids (Thermo Fisher Scientific), 1x Pen/Strep (Thermo Fisher Scientific), and 0.1 mM βME (Thermo Fisher Scientific), maintained at 37°C with 5% CO2.

The wild-type and Xist’s Repeat B deletion female ES cell line (M. musculus/M. castaneus F2 hybrid) harboring a mutated Tsix allele has been previously established ^32,50^. The male ES cell line was previously referred to as J1^50^. ES cells were grown on γ-irradiated mouse embryonic fibroblast (MEF) feeder cells. Culture conditions included DMEM, high glucose, GlutaMAX™ Supplement, pyruvate (Thermo Fisher Scientific), 15% Hyclone FBS (Sigma), 25 mM HEPES pH 7.2-7.5, 1x MEM non-essential amino acids, 1x Pen/Strep, 0.1 mM βME, and 500 U/mL ESGRO recombinant mouse Leukemia Inhibitory Factor (LIF) protein (Sigma) at 37°C with 5% CO_2_.

### ES cell differentiation

Undifferentiated ES cells were initially cultivated on γ-irradiated MEF feeders for 3 days (day 0). Subsequently, the ES cell colonies were trypsinized (Thermo Fisher Scientific), and the feeders were removed. ES cells were then transitioned to a medium devoid of LIF and cultured in suspension for 4 days to form embryoid bodies (EBs). On day 4, the EBs were carefully settled onto gelatin-coated plates and allowed to undergo further differentiation until they were harvested on day 14.

### CHART-seq

Xist CHART-seq methodology was conducted in accordance with a previously described protocol ^51^. Briefly, the cells were collected and suspended in PBS at a concentration of 2.5 million cells/mL. Subsequently, cross-linking was performed using 1% formaldehyde at room temperature for 10 min, followed by quenching with 0.125 M glycine for an additional 10 min. After two washes with ice-cold PBS, cells were pelleted and snap-frozen in liquid nitrogen. For CHART, 25 million cells were thawed on ice and resuspended in 1 mL ice-cold sucrose buffer (10 mM HEPES pH 7.5, 0.3 M sucrose, 1% Triton X-100, 100 mM potassium acetate, 0.1 mM EGTA, 0.5 mM spermidine, 0.15 mM spermine, 1 mM DTT, 1x protease inhibitor cocktail, 10 U/mL SUPERase•In™ RNase Inhibitor). The cell suspension was rotated at 4°C for 10 min, followed by dilution with 2 mL of cold sucrose buffer. Douncing was performed using an RNaseZAP treated 15-mL glass Wheaton Dounce tissue grinder 20 times with a tight pestle. The nuclear suspension was carefully layer on top of a cushion of 7.5 ml glycerol buffer (10 mM HEPES pH 7.5, 25% glycerol, 1 mM EDTA, 0.1 mM EGTA, 100 mM potassium acetate, 0.5 mM spermidine, 0.15 mM spermine, 1 mM DTT, 1x cOmplete EDTA-free protease inhibitor cocktail, and 5 U/mL RNase inhibitor) in a new 15 ml tube, and centrifuged at 1,500 g for 10 min at 4°C. The pellet was resuspended in 3 mL of PBS and cross-linked with 3 ml 6% formaldehyde for 30 min at room temperature with rotation. Afterwards, nuclei were pelleted by centrifugation at 1,000 g for 5 min at 4°C and washed three times with ice-cold PBS, and resuspended in 1 mL ice-cold nuclear extraction buffer (50 mM HEPES, pH 7.5, 250 mM NaCl, 0.1 mM EGTA, 0.5% N-lauroylsarcosine, 0.1% sodium deoxycholate, 5 mM DTT, 10 U/mL RNase inhibitor), rotated for 10 min at 4°C, and then centrifuged at 400 g for 5 min at 4°C and resuspended in 230 μL cold sonication buffer (50 mM HEPES pH 7.5, 75 mM NaCl, 0.1 mM EGTA, 0.5% N-lauroylsarcosine, 0.1% sodium deoxycholate, 0.1% SDS, 5 mM DTT, 10 U/mL RNase inhibitor) to a final volume of ∼270 μL. The nuclei were sonicated in a microtube using a Covaris E220 sonicator (140 W peak incident power, 10% duty factor, 200 cycles/burst, 4°C, 300 s). Sonicated chromatin was centrifuged at 16,000 g for 20 min at 4°C, and transfer the supernatant (∼220 μL) to new tube and add 170 μL sonication buffer to a final volume of ∼390 μL, which was pre-cleared by 60 uL MyOne Streptavidin C1 beads (Thermo Fisher Scientific) in 640 μL 2x hybridization buffer (50 mM Tris pH 7.0, 750 mM NaCl, 1% SDS, 1 mM EDTA, 15% formamide, 1 mM DTT, 1 mM PMSF, 1x cOmplete EDTA-free protease inhibitor cocktail, 100 U/mL RNase inhibitor) at room temperature for 1 h with rotation. Pre-cleared chromatin was divided into two CHART reactions for Xist and control capture, and 1% was saved as an input sample. For each CHART reaction, 36 pmol of antisense (Xist) or sense (control) biotinylated capture probes (pooled probe as previously described) were used ^21^. Hybridization was performed at room temperature with rotation overnight, and 120 μL C1 beads were added to the samples and incubated at 37°C for 1 h with rotation. The beads were washed once with 1x hybridization buffer (33% sonication buffer, 67% 2x hybridization buffer) at 37°C for 10 min, five times with wash buffer (10 mM HEPES pH 7.5, 150 mM NaCl, 2% SDS, 2 mM EDTA, 2 mM EGTA, 1 mM DTT) at 37°C for 5 min, and twice with elution buffer (10 mM HEPES pH 7.5, 150 mM NaCl, 0.5% NP-40, 3 mM MgCl2, 10 mM DTT) at 37°C for 5 min. 1% of the final wash was saved as an “RNA pulldown” sample. CHART-enriched DNA was eluted twice in 200 μL of elution buffer supplemented with 5 U/μL RNase H (New England BioLabs) at room temperature for 20 min. The “input” sample and CHART DNA were treated with 0.5 mg/mL RNase A (Thermo Fisher Scientific) at 37°C for 1 hr with rotation and then incubated with 1% SDS, 10 mM EDTA, and 0.5 mg/mL proteinase K (Sigma) at 55°C for 1 hr. Reverse crosslinking was performed using 150 mM NaCl (final concentration 300 mM) at 65°C overnight. DNA was purified using phenol-chloroform and further sheared to ∼300bp fragments using Covaris E220e (140 W peak incident power, 10% duty factor, 200 cycles/burst, 120 s, 4°C). Sonicated DNA was purified using 1.8x Agencourt AMPure XP beads (Beckman Coulter). Input and CHART DNA libraries were prepared according to the protocol of the NEBNext® Ultra™ II DNA Library Prep Kit for Illumina (NEB E7645S). Libraries were sequenced on Novaseq S4, generating approximately 30 million 150-nt paired-end reads per sample.

### Data source and processing

Raw data for CHART-seq and bulk RNA-seq generated in this study have been deposited in the Gene Expression Omnibus (GEO) database with accession number GSE271096 and GSE271097, respectively. ChIP-seq data utilized in this study were sourced from a study conducted by David et al. in 2020 ^33^ with GEO accession number: GSE135389. Additionally, RNA-seq and ChIP-seq data specifically pertaining to Xist inhibitor X1 were obtained from a dataset published in 2022 ^36^ with GEO accession number: GSE141683. RNA-seq data for Xist TG in mES cells were acquired from another dataset published in 2017 ^35^ with GEO accession number: GSE92894. RNA-seq data for male and female MEF cells were acquired from two published dataset with GEO accession number: GSE246699 and GSE118443. The data were processed and re-analyzed consistently in this study.

### CHART-seq and ChIP-seq analysis

Initial data preprocessing involved removing adaptors using trim_galore/cutadapt (versions 0.4.3/1.7.1). Subsequently, trimmed reads were prepared for alignment. Alignment of the genomes of Mus musculus (mus) and Mus castaneus (cas) was performed using NovoAlign (version 4.03). Aligned reads were then mapped back to the reference mm10 genome using SNPs to generate a BAM file ^28^. Samtools (version 1.11) was employed for random sampling to ensure uniform library sizes across the samples. Utilizing DeepTools (version 3.1.2), input-subtracted ChIP and CHART coverage profiles were created, resulting in bigwig. To discern the peaks of Xist CHART on the autosomes, chrX reads were initially excluded from the BAM file. MACS2 (version 2.1) was used for peak calling using default parameters. The resultant peak files from two replicates were merged using bedtools (version 2.30) intersect. Bedtools window (version 2.30) facilitated the creation of gene lists based on peak sizes, spanning 10, 20, 50, and 100 kilobases. Deeptools was used to conduct comparative analyses of coverage, replicate correlation analysis, represented through either profile or heatmap plots, for genomic regions associated with Xist, H3K27me3, or H2A119ub across distinct categories of gene lists. IDR (Irreproducible Discovery Rate, version 2.0.2) was used to check the reproducibility of peaks identified in replicates. Intervene (version 6.0.2) was used to generate the Venn diagram of the peak.

### RNA-seq

Total RNA was extracted using TRIzol (Thermo Fisher Scientific), and rRNA was selectively removed using the NEBNext rRNA Depletion Kit (New England BioLabs), according to the manufacturer’s instructions. RNA-seq libraries were prepared using the NEBNext Ultra II Directional RNA Library Prep Kit for Illumina (New England BioLabs), which enabled the creation of strand-specific RNA-seq libraries. Libraries were sequenced on Novaseq S4, generating ∼30 million 150-nt paired-end reads per sample.

### RNA-seq analysis

Raw RNA-seq data underwent adaptor removal and trimming using trim_galore/cutadapt (versions 0.4.3/1.7.1). RNA-seq data were mapped to three different genomes: C57BL/6J (mm10), M. musculus (mus), and M. castaneus (cas). By employing Subread (version 2.0.2) featureCounts ^52^, counts per gene were computed, while deeptools were utilized to generate bigwig files, providing a comprehensive overview of read coverage across the genomes. For expression quantification, both Reads Per Kilobase Million (RPKM) and log transformed RPKM values were calculated to establish a robust foundation for downstream analysis.

### Quantification and Statistical Analysis

In the experimental design, two replicates were used for both the CHART and RNA-seq analyses. Statistical analyses were conducted, and the corresponding p-values are transparently reported in the figures and their respective legends. The significance levels are denoted by asterisks, where *, **, ***, and **** represent p < 0.05, p < 0.01, p < 0.001, and p < 0.0001, respectively.

## ACKNOWLEDGMENTS

We acknowledge all members of the Lee lab for their invaluable contributions, insightful comments, and stimulating discussions. We thank Dr. Uri Weissbein for his assistance with the CHART protocol. This work was supported by an NIH grant (R01-HD097665) to JTL.

## SUPPLEMENTAL TABLE LEGENDS

Table S1-S3. Xist binding peaks on autosomal regions (MACS2 peak calling) information at day4 (S1), day7 (S2), and day14 (S3) in WT female mouse ES cells.

Table S4-S6. Xist target genes on autosomal regions (10kb among the binding peak) information at day4 (S4), day7 (S5), and day14 (S6) in WT female mouse ES cells.

Table S7-S9. Xist binding peaks on autosomal regions (MACS2 peak calling) information on WT (S7), ΔRepB (S8), and ΔRepE (S9) female MEFs.

Table S10. Xist target genes on autosomal regions (10kb among the binding peak) information on WT female MEFs.

Table S11. Statistical information in this manuscript.

---

**Figure S1.**
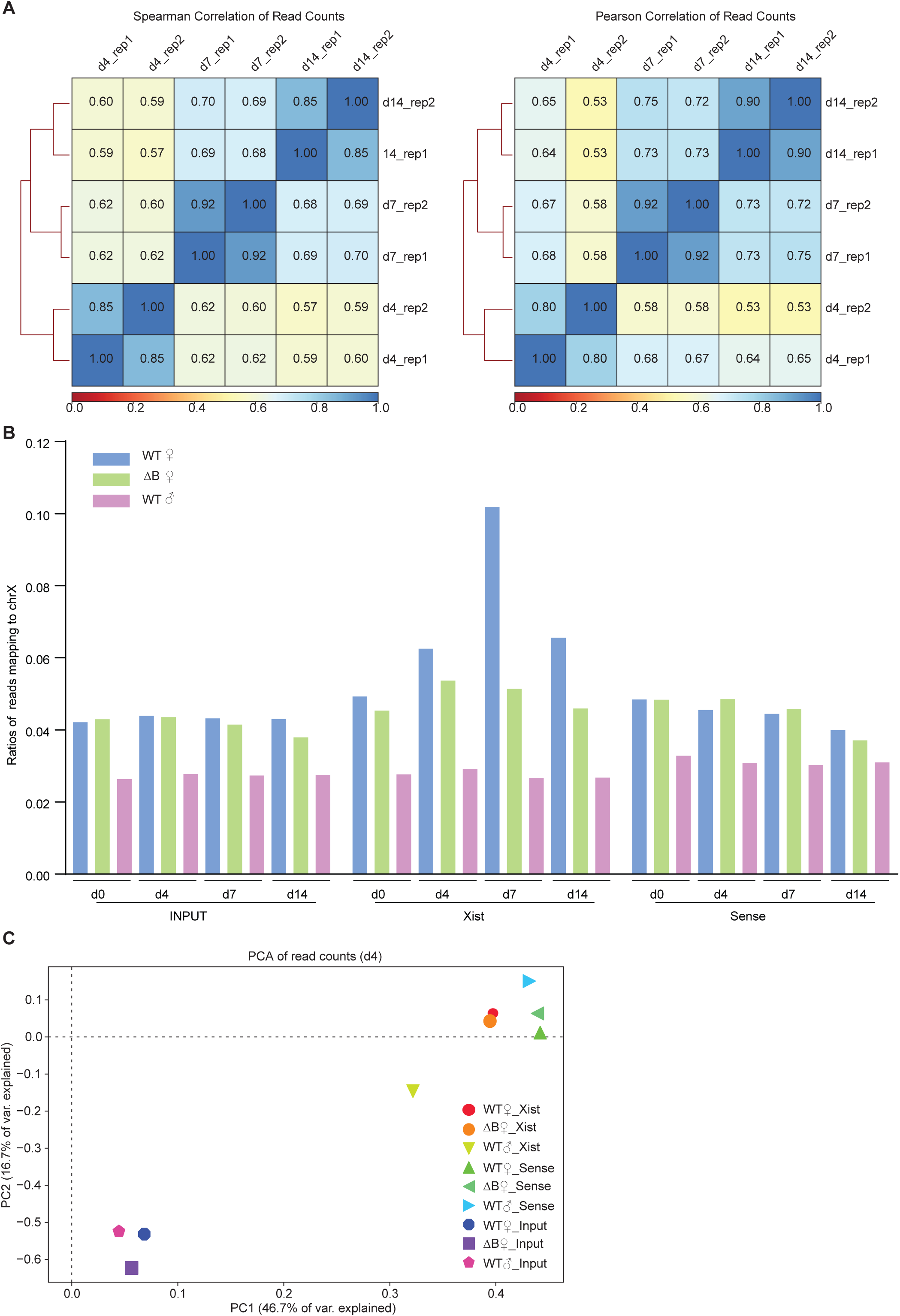
Xist binds autosomal genes in trans. **A.** Spearman (rank-based relationship) and Pearson (linear relationship) correlation analysis show that a strong positive correlation between the two biological CHART sequencing. **B.** Xist CHART-seq reads percent of autosomes and Chromosome X in WT and ΔRepB female ES cells at day 0, day 4, day 7, and day 14. ES day 7 exhibits the highest Xist coverage. Sense probe and male ES cells are used as a control. **C.** Principal Component Analysis (PCA) of CHART-seq reads include Xist, sense, and input in WT and ΔRepB female, and male ES cells at day 4. Samples clustering closer together share similar genome-wide coverage patterns.

**Figure S2.**
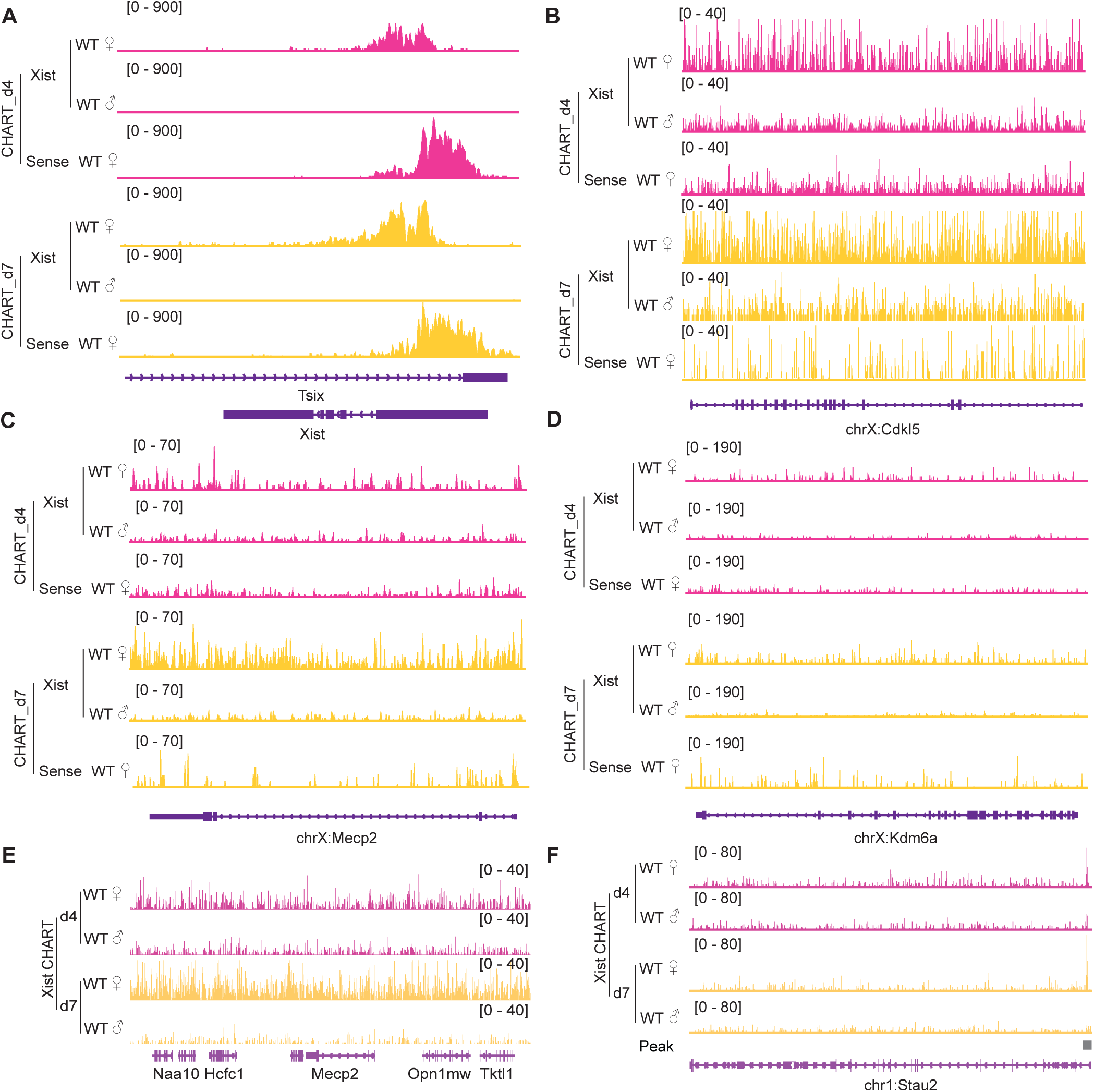
Xist binds X chromosome during female ES cell differentiation. **A-D.** Representative binding signals of Xist RNA on Xist locus (A), genes subjective to XCI such as *Cdkl5* (B), *Mecp2* (C), and escapee genes (*Kdm6a*) (D) in WT female ES cells at day 4 and day 7. WT male ES cells and sense probe are used as control. **E.** Representative consecutive binding signals of Xist on chromosome X (∼320 kb) in WT female ES cells at day 4 and day 7. WT male ES cells are used as control. **F.** Representative site-specific binding of Xist on autosome locus (*Stau2*) in WT female ES cells at day 4 and day 7, WT male ES cells are used as control.

**Figure S3.**
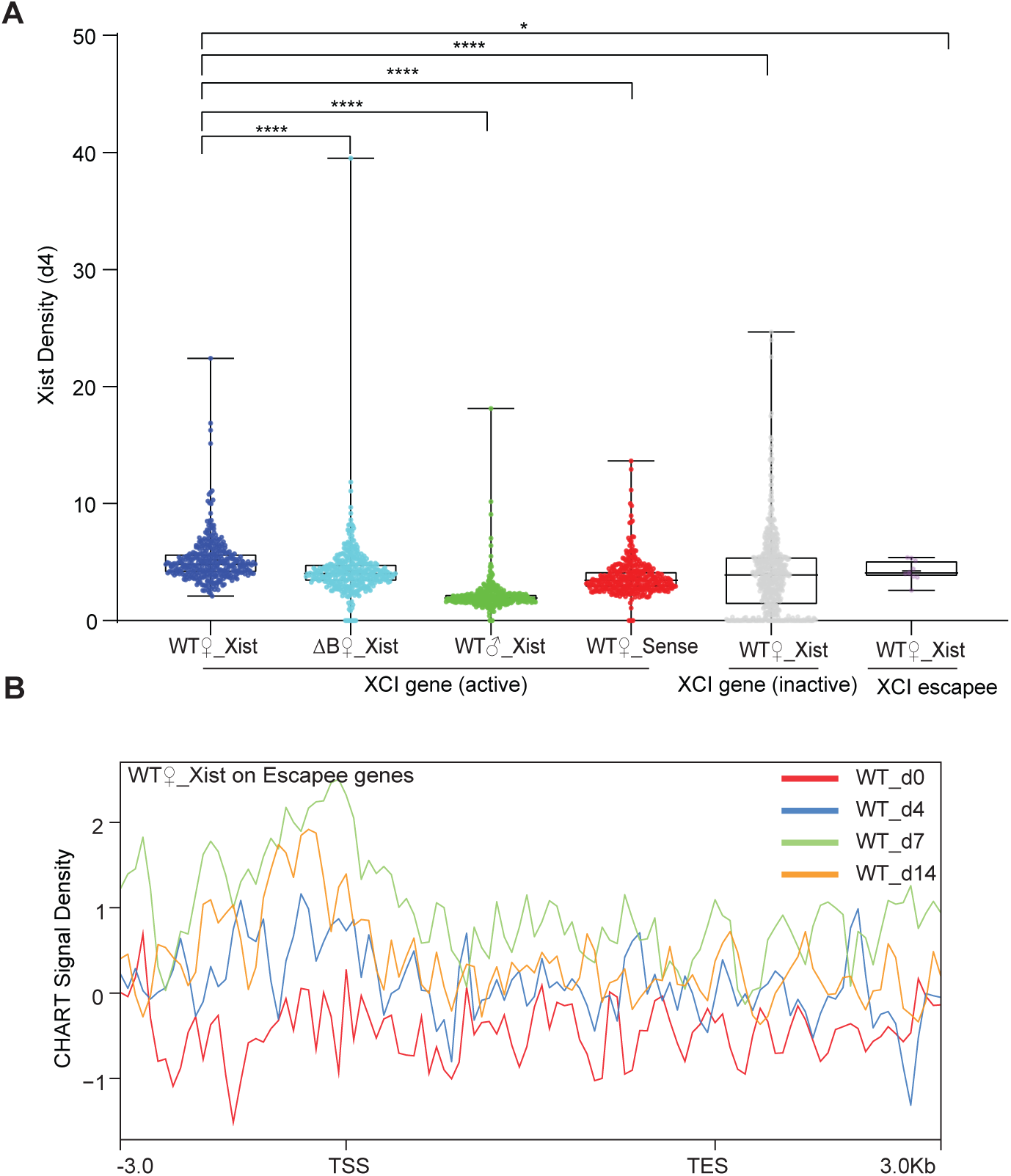
Xist binds XCI genes during differentiation. **A.** Xist CHART signal coverage on X chromosome genes (XCI-active/inactive, and escapee) in WT female ES cells at day 4. Sense and male ES cells are used as control. P-values are determined using the Wilcoxon rank sum test. **B.** Average profile of Xist CHART signal on XCI escapee genes (day 0, 4, 7, and 14) shows that Xist accumulated in the upstream of the promoter but depleted in the escapee gene body.

**Figure S4.**
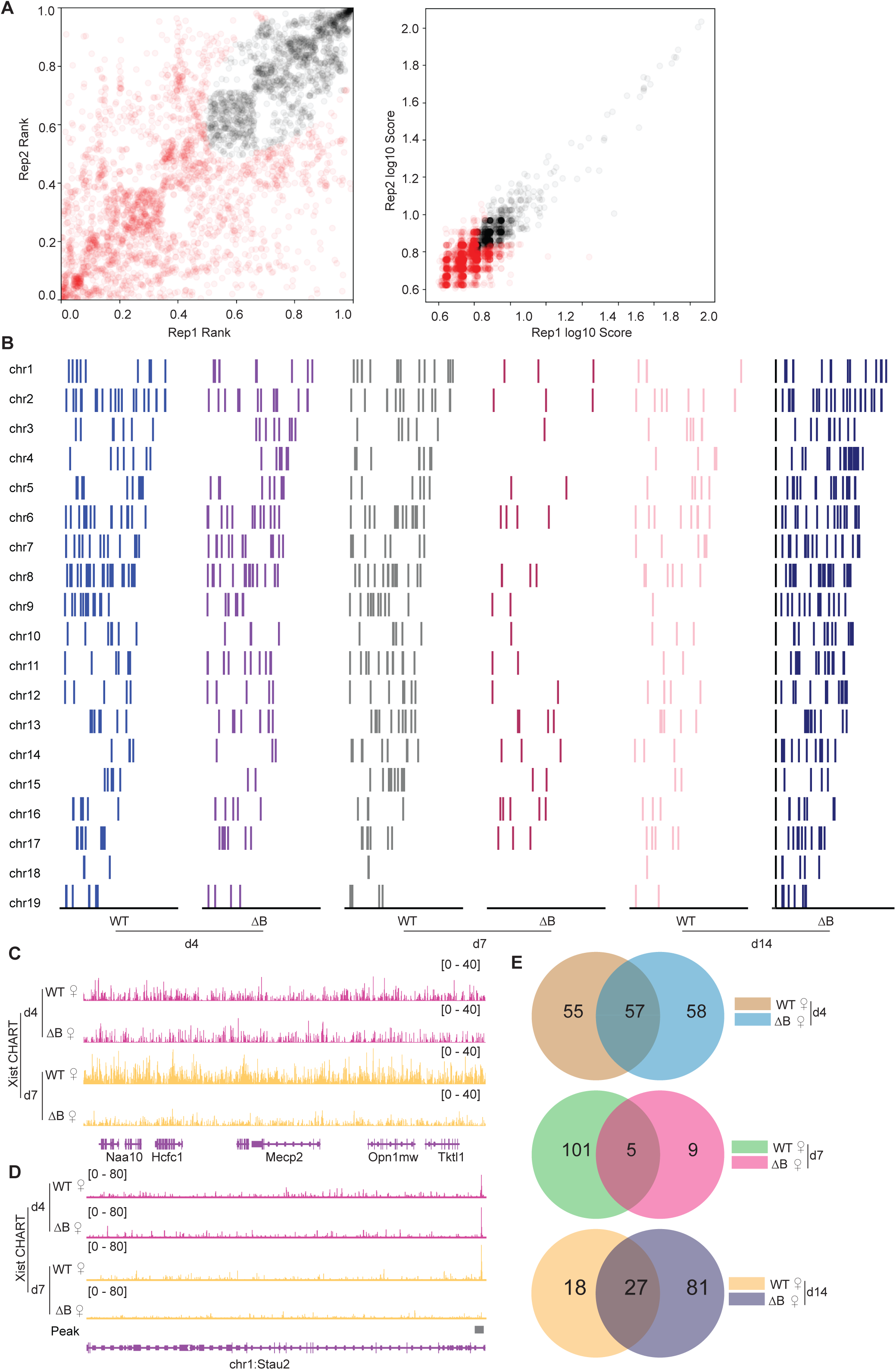
Xist binds select autosomal genes in trans. **A.** IDR (Irreproducible Discovery Rate) analysis to assess the reproducibility of the peaks detected across biological replicates (day 4). The results showed a strong correlation between the replicates, with an IDR threshold of 0.05 (red point > 0.05). **B.** Xist peak pattern (MACS2 peak calling) on autosomes in WT and ΔRepB female ES cells at day 4, day 7, and day 14. **C.** Representative consecutive binding signals of Xist on chromosome X (∼320 kb) in WT and ΔRepB female ES cells at day 4 and day 7. **D.** Representative site-specific binding of Xist on autosome locus (*Stau2*) in WT and ΔRepB female ES cells at day 4 and day 7. **E.** The Venn diagram illustrates the overlap of peak sites identified in WT and ΔRepB female ES cells at day 4, day 7, and day 14.

**Figure S5.**
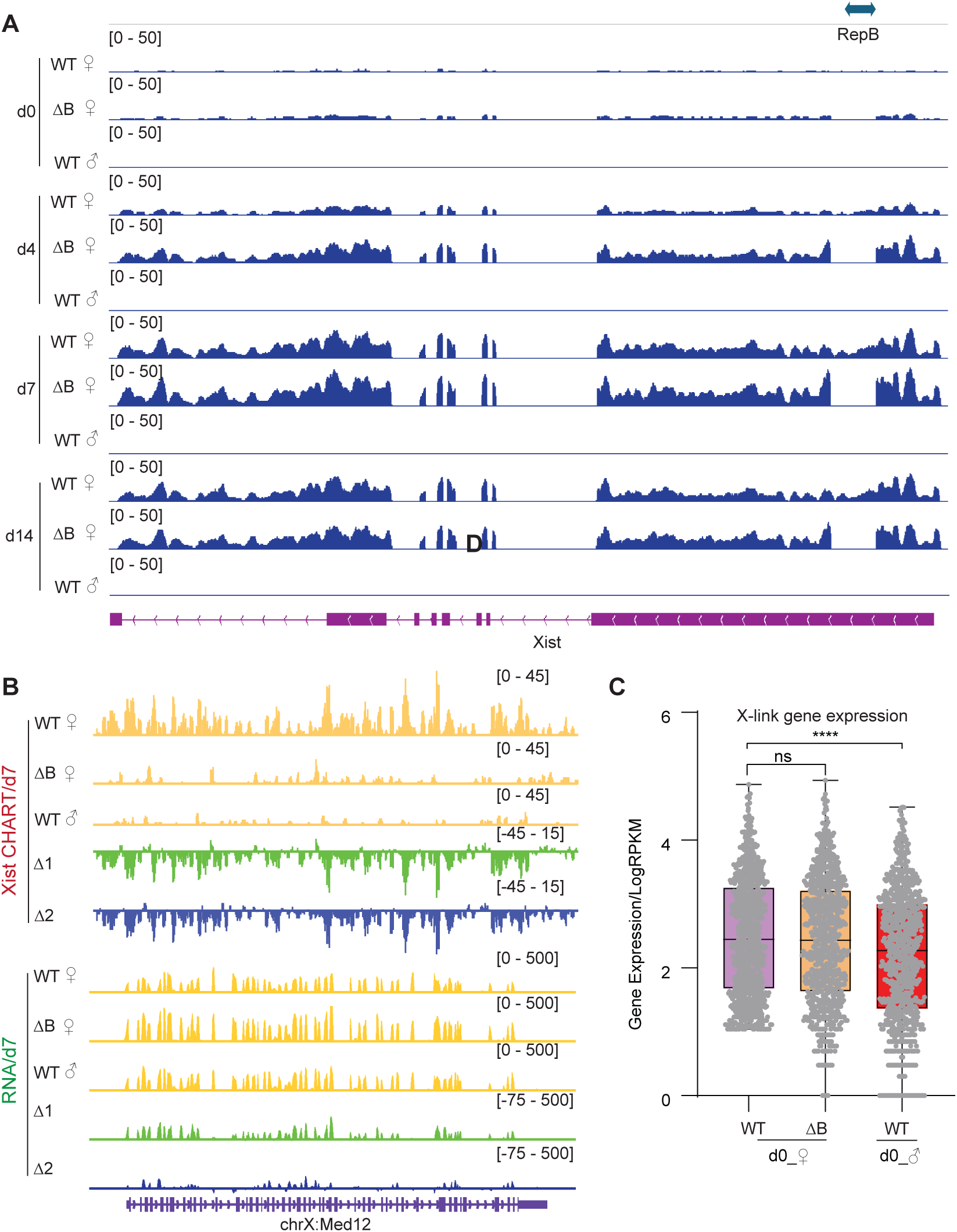
Xist binds X-linked genes is Xist’s RepB dependent. **A.** RNA-seq track patterns of Xist in WT, ΔRepB female, and male ES cells at day 0, day 4, day 7, and day 14. **B.** Exemplifying CHART-seq and RNA-Seq patterns of an X-linked gene (*Med12*) at day 7. Change in coverage (Δ1 and Δ2) is shown below (Δ1 for ΔRepB♀ -WT♀, and Δ2 for WT♂ -WT♀). **C.** Evaluation of gene expression levels for X-linked genes in WT and ΔRepB female, and male ES cells at day 0. P-values are determined using the Wilcoxon rank sum test.

**Figure S6.**
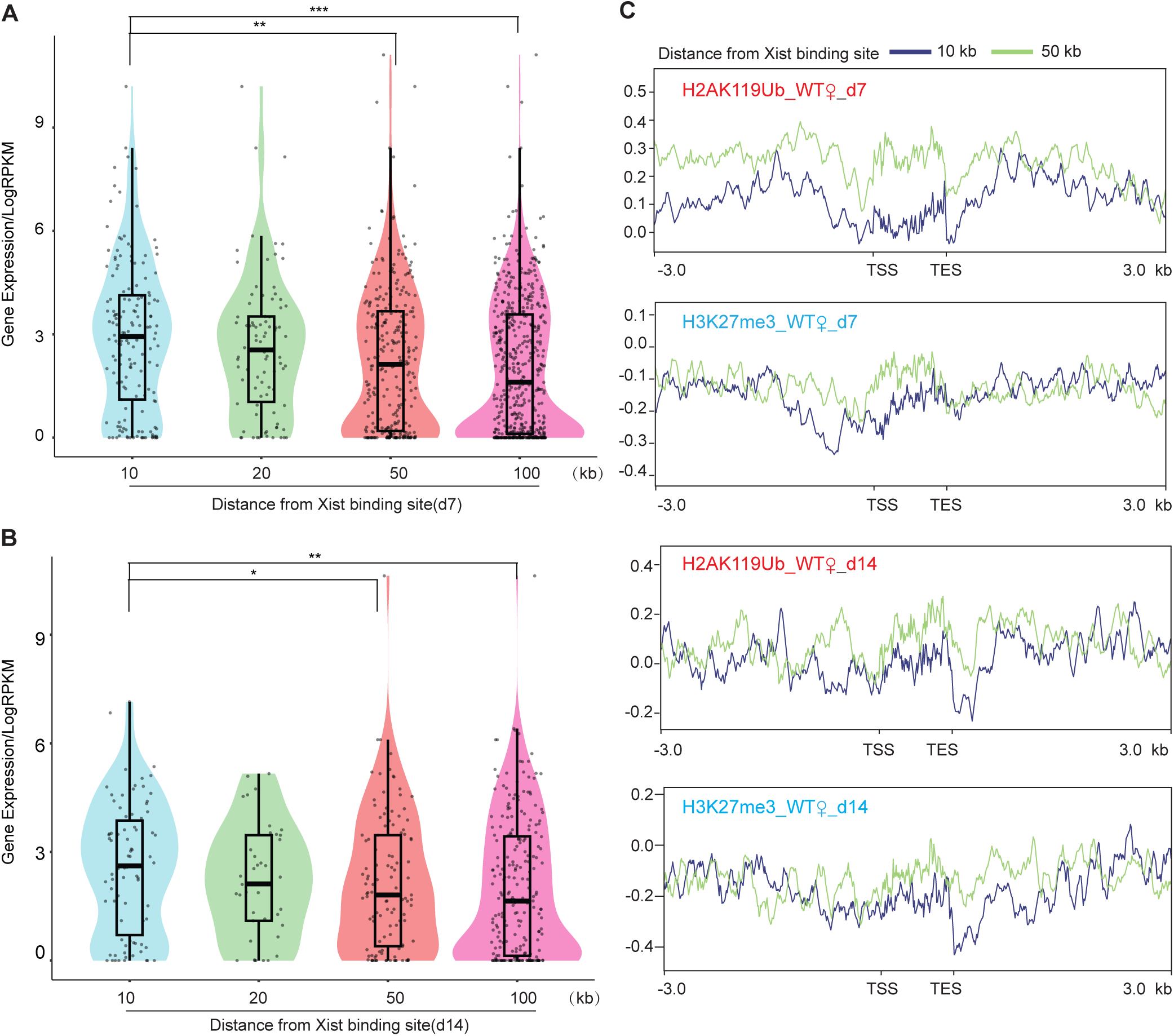
Genes bound by Xist exhibit higher expression levels and lower H3K27me3 and H2AK119ub binding levels. **A-B.** The analysis of gene expression within the 10, 20, 50, and 100-kilobase binding regions of Xist is performed in WT female ES cells at day 7 (B), and day 14 (C), respectively. P-values are determined using the Wilcoxon rank sum test. **C.** Average profile plots showing H3K27me3 and H2AK119ub coverage over genes within the 10 and 50-kilobase binding regions of Xist in WT female ES cells.

**Figure S7.**
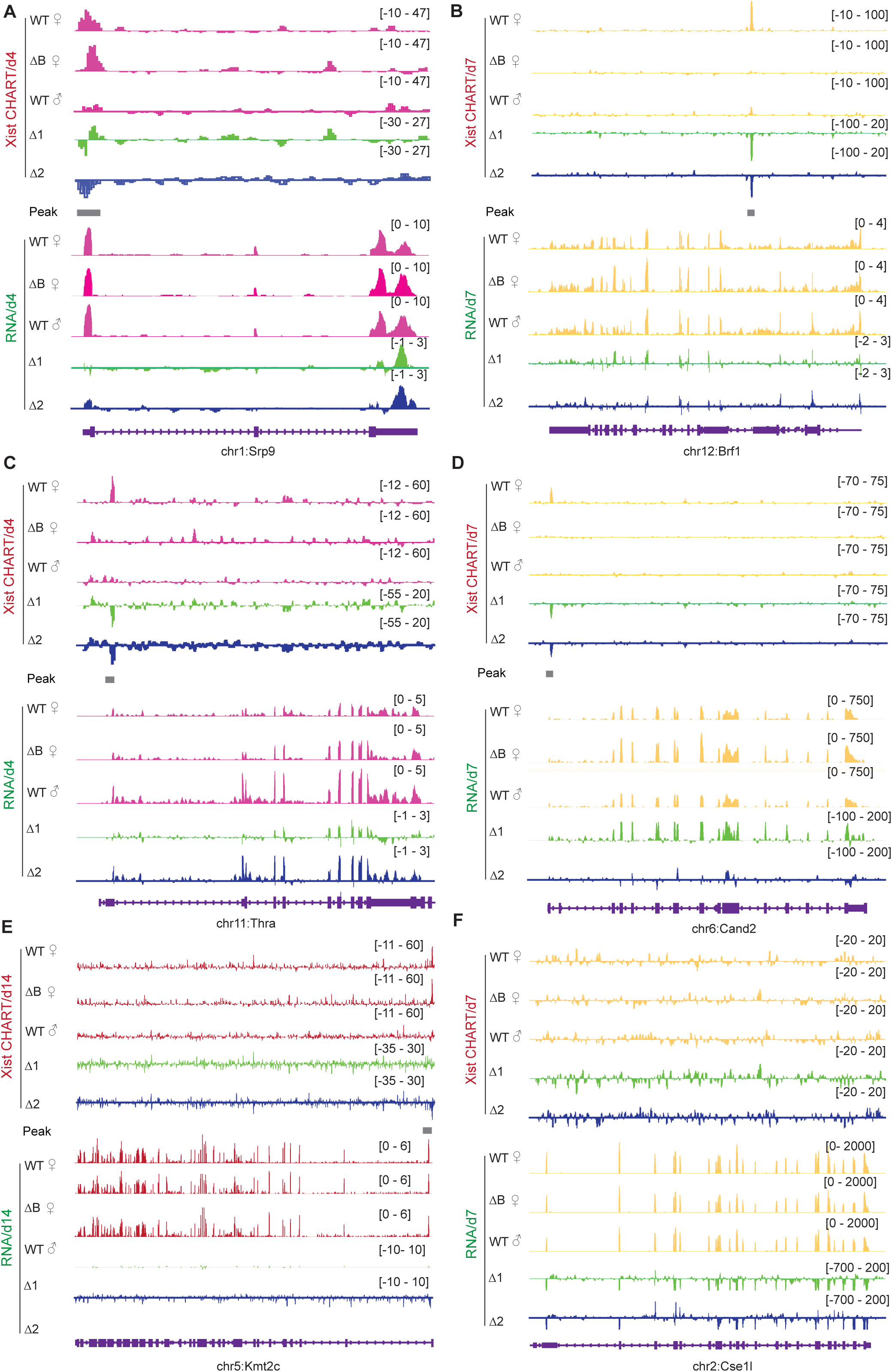
Example of Xist binding on autosomal genes and influence on gene expression. This figure illustrates CHART-seq and RNA-seq patterns of autosomal genes, including *Srp9, Brf1, Thra, Cand2, and Kmt2c*, which exhibit Xist binding on different days. *Ces1l*, which lacks Xist binding, is used as a control. Change in coverage (Δ1 and Δ2) is shown below (Δ1 for ΔRepB♀ -WT♀, and Δ2 for WT♂ -WT♀).

**Figure S8.**
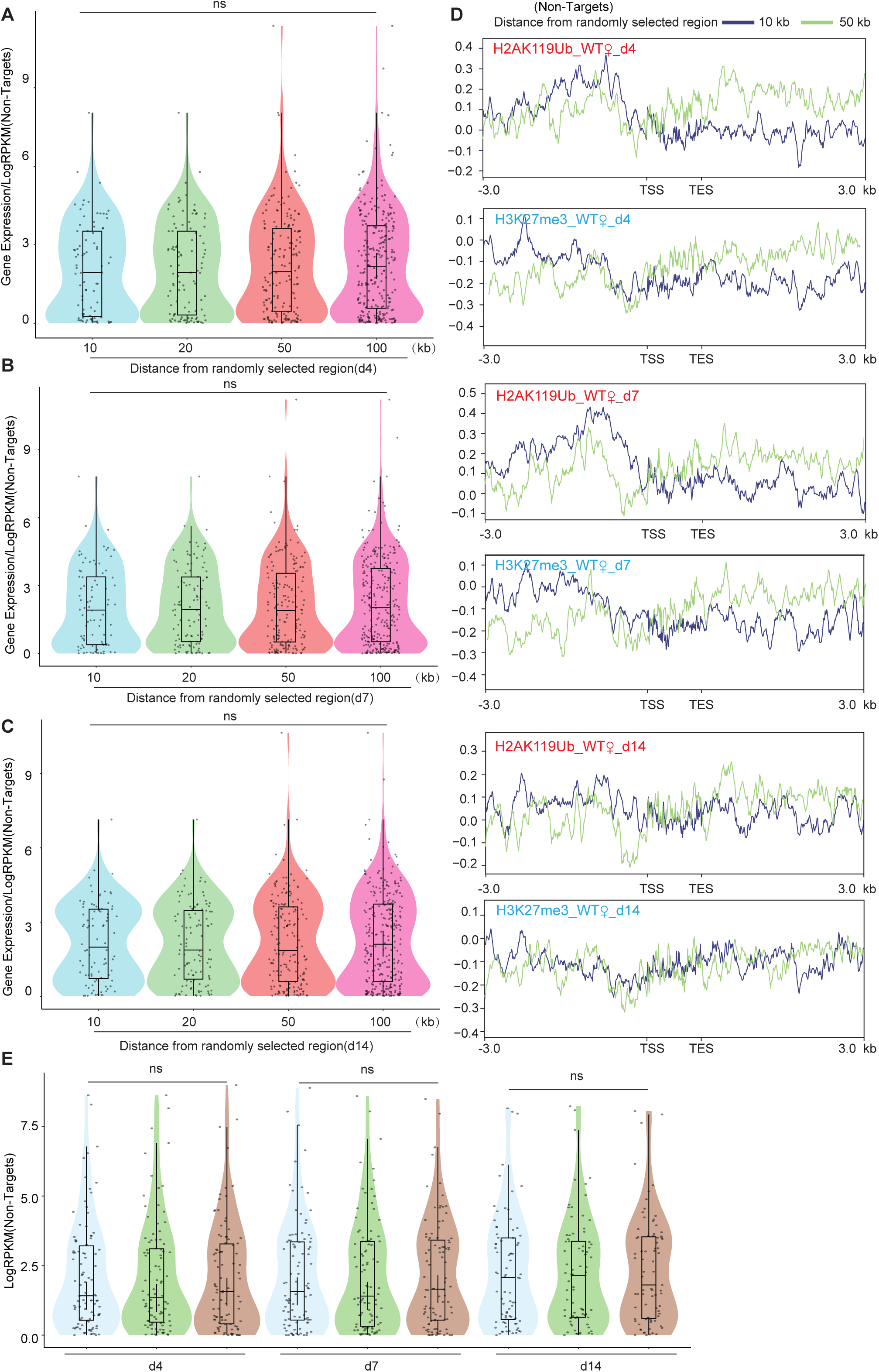
Genes not bound by Xist exhibit no changes in gene expression or differences in H3K27me3 and H2AK119ub signals. **A-C**. The analysis of gene expression within the 10, 20, 50, and 100-kilobase randomly selected regions is performed in WT female ES cells at day 4 (A), day 7 (B), and day 14 (C), respectively. P-values are determined using the Wilcoxon rank sum test. **D.** Average profile plots showing H3K27me3 and H2AK119ub coverage over genes within the 10 and 50-kilobase randomly selected regions in WT female ES cells at different time points. **E.** Assessing gene expression levels of Xist non-targets on autosomes in WT, ΔRepB female ES cells, and male ES cells at different time points. P-values are determined using the Wilcoxon rank sum test.

**Figure S9.**
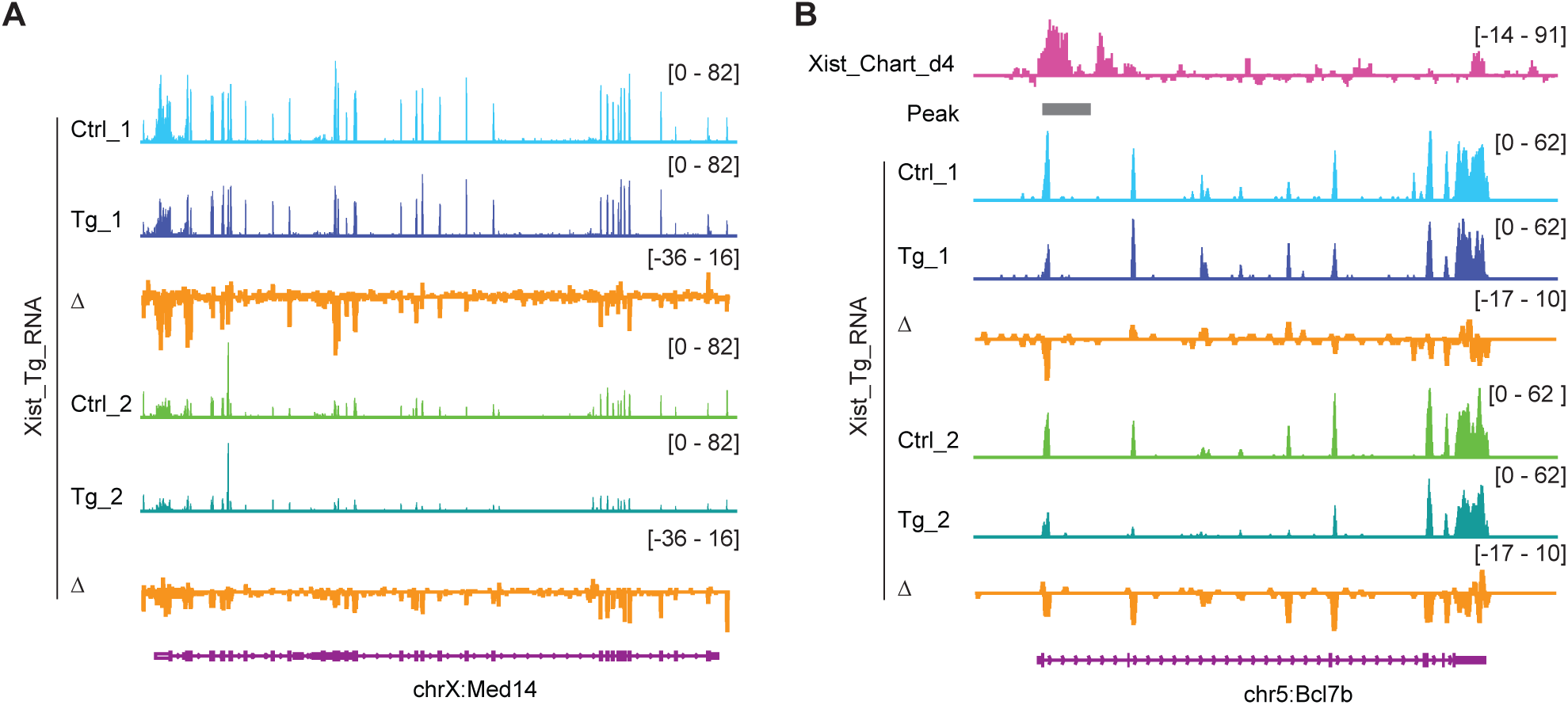
Xist overexpression inhibits select autosomal genes and X-linked genes. RNA-seq track shows the *Med14* (X-linked gene) (A) and *Bcl7b* (an Xist autosomal target gene) (B) expression levels in differentiated ectopic Xist overexpressed ES cell lines (Tg), the control (Ctrl) was doxycycline-treated wildtype cell. Change in coverage (Δ) is shown below (Tg - Ctrl).

**Figure S10.**
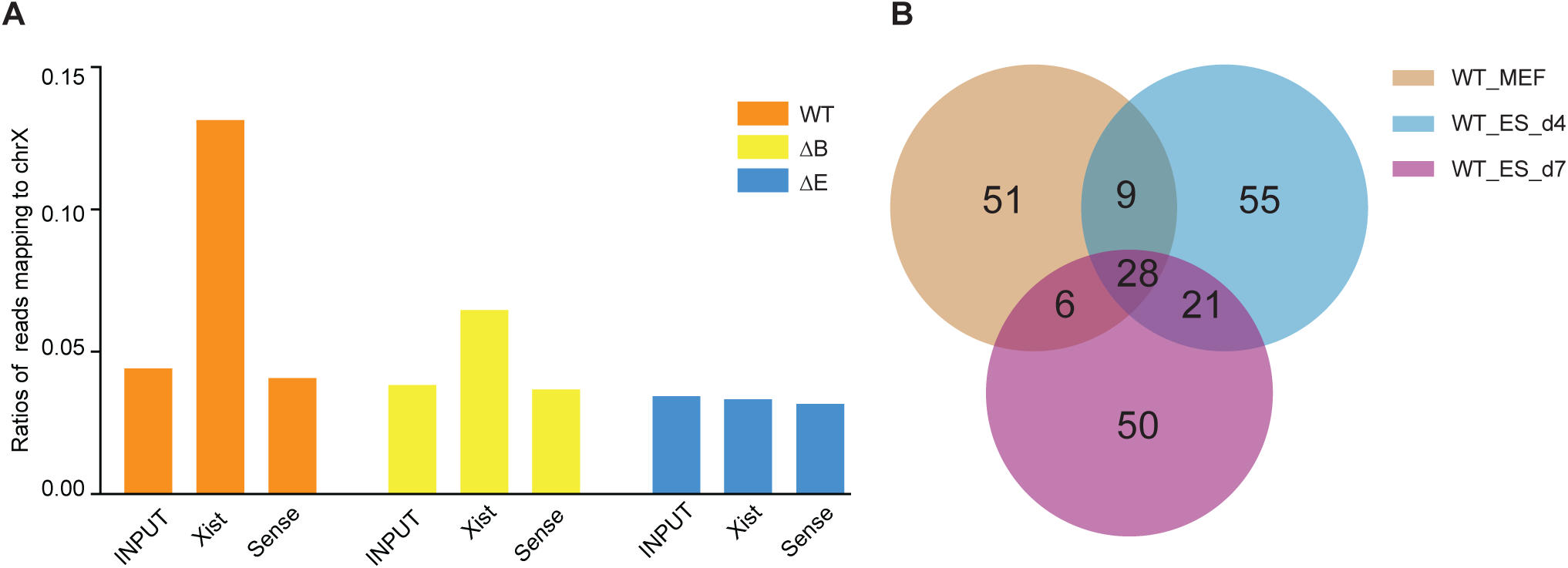
Xist has overlapped autosomal binding peaks in differentiated ES and MEF cells. **A.** Coverage of CHART-Seq reads (input, Xist, and sense control) on Chromosome X in WT, ΔRepB, and ΔRepE female MEF cells. **B.** The Venn diagram illustrates the overlap of peak sites identified in WT female ES cells at day 4, day 7 and MEF cells.

